# Structure-informed functional connectivity driven by identifiable and state-specific control regions

**DOI:** 10.1101/2020.07.10.197046

**Authors:** Benjamin Chiêm, Frédéric Crevecoeur, Jean-Charles Delvenne

## Abstract

A challenge in neuroscience is to describe the contribution of the brain anatomical wiring to the emergence of coordinated neural activity underlying complex behavior. Indeed, patterns of remote coactivations that adjust with the ongoing task-demand do not systematically match direct, static anatomical links. Here, we propose that observed coactivation patterns, known as Functional Connectivity (FC), can be explained by a linear diffusion dynamics defined on the brain architecture and driven by control regions. Our model, termed *structure-informed* FC, is based on a novel interpretation of functional connectivity according to which different sets of brain regions controlling the information flow on a fixed anatomical wiring enable the emergence of state-specific FC. This observation leads us to introduce a framework for the identification of potential control centers in the brain. We find that well-defined, sparse and robust sets of control regions, which partially overlap across several task conditions and resting-state, produce FC patterns comparable to empirical ones. In conclusion, this work introduces a principled method for identifying brain regions underlying the task-specific control of brain activity.

**Significance statement:** Understanding how brain anatomy promotes particular patterns of coactivations among neural regions is a key challenge in neuroscience. This challenge can be addressed using network science and systems theory. Here, we propose that coactivations result from the diffusion of information through the network of anatomical links connecting brain regions, with certain regions controlling the dynamics. We translate this hypothesis into a model called *structure-informed functional connectivity*, and we introduce a framework for identifying control regions based on empirical data. We find that our model produces coactivation patterns comparable to empirical ones, and that distinct sets of control regions are associated with different functional states. These findings suggest that controllability is an important feature allowing the brain to reach different states.

## Introduction

Recently, approaches combining Magnetic Resonance Imaging (MRI) and network science emerged in order to characterize links among neural Regions of Interest (ROIs) [1, 2]. Most studies focus either on structural connections or on functional interactions, which capture two distinct aspects of brain connectivity. On the one hand, diffusion MRI (dMRI) with tractography [3] enables the mapping of white matter pathways and describes the anatomical links between ROIs. This structural description of the human brain forms a network called the *connectome* [4, 5]. On the other hand, the Blood-Oxygenation-Level Dependent (BOLD) signal in functional MRI (fMRI) provides an estimate of brain activity in grey matter areas [6]. The matrix of pairwise Pearson’s correlation coefficients between regional BOLD time series is a common tool to quantify *functional connectivity* (FC) among ROIs [1, 2]. Unlike the connectome, functional connectivity varies over short timescales and across resting-state and task conditions [7].

An important challenge in neuroscience is to characterize the relationship between the connectome and functional connectivity [8, 9, 10, 11]. Several approaches have been proposed in the literature in order to elucidate this relationship in macroscale brain networks and understand the importance of the anatomical organization in promoting particular patterns of activity. Along with methods based on graph signal processing and spectral decompositions [12, 13], it has been proposed that describing the link between the connectome and FC requires a model of information flow between ROIs [14]. For instance, models based on random walks and diffusion on the connectome have been able to partly reproduce resting-state FC [15, 16, 17]. Viewing the brain as a dynamical system allows us to study the controllability of this system, i.e. its ability to account for context-dependent control signals in order to affect the overall state of the brain. [18, 19, 20]. The framework of network controllability requires to define input regions, i.e. ROIs capable of integrating control signals [21]. Earlier work demonstrated that any single input region was theoretically sufficient to get controllability of the connectome [18, 22, 23]. One shortcoming is that although controllable in theory, some configurations are practically unfeasible as they would require excessive control energy. Moreover, several pieces of evidence from the fields of motor and cognitive control suggest that sets of regions are responsible for the control of brain activity [24, 25, 7, 26, 27]. Despite these advances in brain communication modelling and connectome controllability, an integrated explanation for the emergence of multiple FC patterns from the static structure of the connectome is still lacking.

Here, we develop a principled approach modelling state-specific FC on the connectome. We leverage the observation that the Gramian matrix used in controllability studies [28, 18] corresponds to the covariance matrix of the activities in the different nodes of a network, assuming a linear transition dynamics among them. This observation brings us to introduce the concept of *structure-informed* FC, i.e. the pairwise functional correlation matrix derived from the structure of the connectome. Since this matrix depends on the choice of input nodes, we show that it is possible to identify the set of input ROIs maximizing the mapping between structure-informed and empirical FC in different states. Using dMRI and fMRI data (resting-state and 7 tasks) from the Human Connectome Project [29], we find that sparse input sets produce FC matrices that are comparable to empirical ones. Moreover, we show that the identified sets are well defined, stable, and state-specific. We discuss their properties and the fact that the method is able to capture the singularity of resting-state compared to the other task-related conditions. Overall, our approach relies on a model linking structure and function in brain networks in order to identify possible subsets of brain regions underlying task-specific control.

## Results

### Structure-informed functional connectivity

In order to investigate how the connectome shapes Functional Connectivity (FC), we study the covariance matrix of a linear dynamics defined on the connectome. In a network of *n* nodes, let **x**(*k*) be the *n*-dimensional state-vector containing the activity level of each node at time *k*. The trajectory of **x** is governed by the following equation:

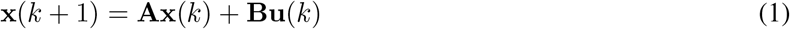

Here, the *n* × *n* system matrix **A** describes the interactions among the nodes of the network, the columns of the *n* × *m* input matrix **B** are canonical vectors identifying the *m* input nodes and **u**(*k*) is an *m*-dimensional vector providing the value of external input signals at time *k*.

When the inputs to the system, i.e. the signals in **u**, are white noise signals, it can be shown that the steady-state covariance matrix of the states, **Σ**= Cov(**x**) satisfies the following Lyapunov equation (see Methods for the derivation):

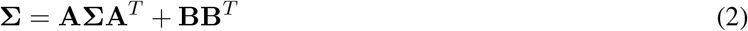

Here, we see that the solution **Σ** depends on the structure of the network and the dynamical model through the system matrix **A**, and on the set of input nodes defined by **B** (Figure 1A). The solution to Equation (2) is known as the controllability Gramian. Here, in contrast to previous studies where **Σ** is used to derive quantitative control properties of individual nodes in the network [28, 18, 30], we interpret the Gramian as the state-covariance matrix obtained by stochastic excitation of the system through a set of control nodes. This allows us to relate it to the concept of functional connectivity. Indeed, after variance normalization, **Σ** becomes a correlation matrix 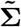 (see Methods) and constitutes the FC matrix associated with the network and its dynamics, which we term the *Structure-Informed* FC and denote **F**_*SI*_:

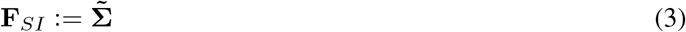

**Figure 1:**
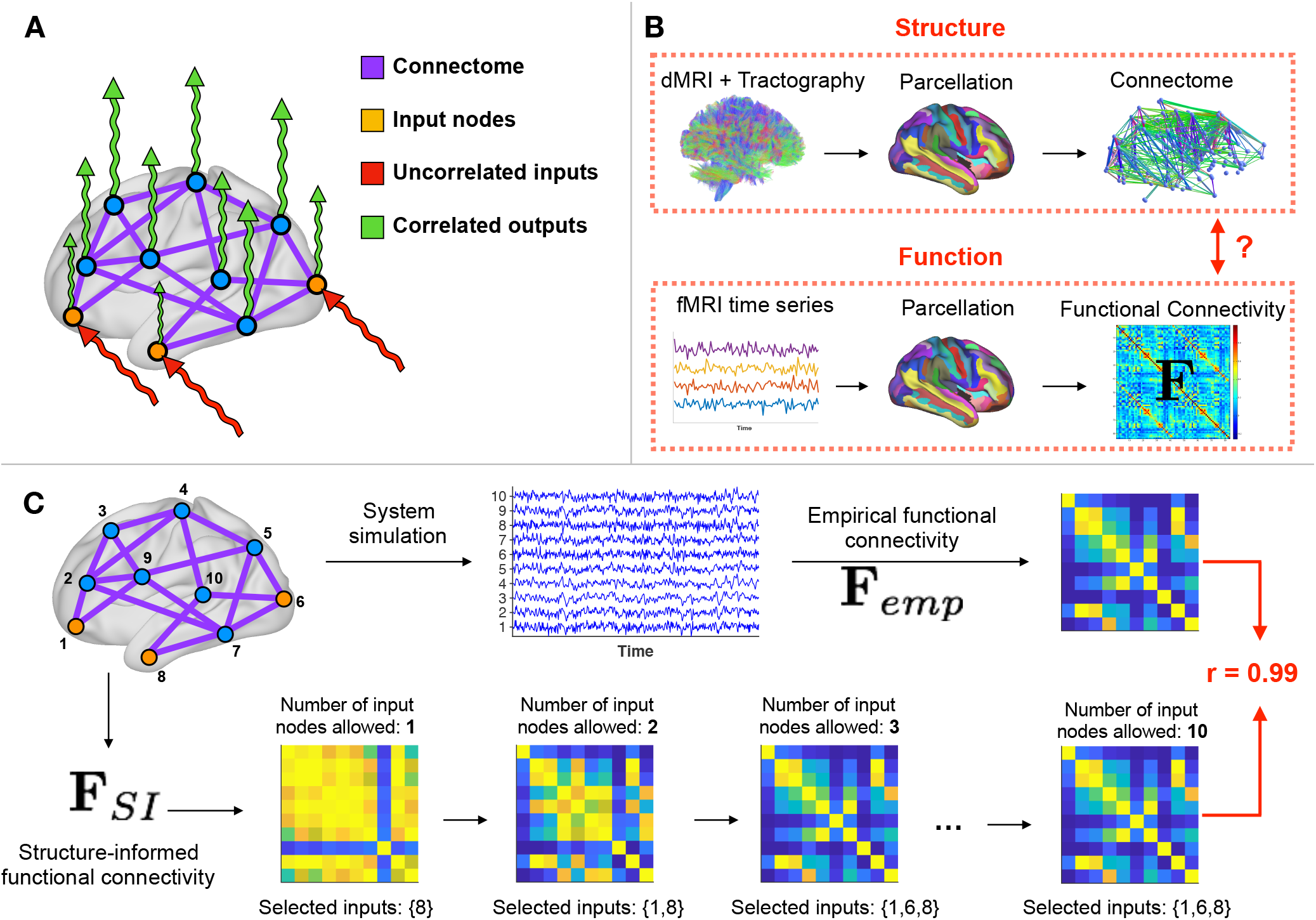
Overview of the approach. **A)** In order to investigate how the connectome shapes functional connectivity, we define a diffusion dynamics on the connectome (purple) and excite it with uncorrelated signals (white noise, red). Depending on the set of input nodes (orange) driving the dynamics, the output signals (green) present correlations patterns that are similar to empirical data. **B)** Data processing workflow. (Top row) We extract the connectome using diffusion imaging (dMRI) and tractography. Nodes correspond to Regions of Interest (ROIs) from a predefined automatic parcellation. (Bottom row) At each ROI, we also retrieve the fMRI BOLD time serie and compute the functional connectivity matrix **F**_*emp*_ between these signals. This step is repeated for 7 tasks and resting-state. **C)** Example on simulated data. We start from a network of *n* = 10 nodes, with uniformly distributed random weights on the edges. (Top row) We choose a set of *m* = 3 input nodes, simulate the noise diffusion process and compute the empirical functional connectivity matrix **F**_*emp*_. (Bottom row) Our framework applied to the network identifies the correct set of input nodes and generates a structure-informed functional connectivity matrix **F**_*SI*_ comparable to the empirical one.

Using the mathematical relation between the network structure and the correlation matrix of the system, we turn to the problem of identifying the set of control inputs defined by **B**, given an empirical FC matrix **F**_*emp*_ obtained from external recordings of the system. For that, we formulate the optimization problem

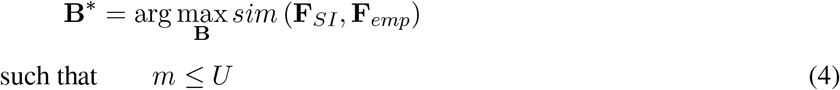

where **F**_*SI*_ is a function of **B**, *m* is the number of columns of **B**, i.e. the cardinality of the input set, and *U* is an upper bound to be fixed in order to control the number of input nodes.

In the present work, we apply this approach to connectomes and FC matrices extracted from MRI data (Figure 1B). We consider a diffusion dynamics to model interactions among ROIs in the connectome (see Methods) as suggested in previous studies on large-scale brain communication [15, 14]. The similarity between **F**_*SI*_ and **F**_*emp*_ is computed as the entry-wise Pearson’s correlation coefficient *r*, following previous work [15, 16, 17, 31, 32]. We refer to this measure of similarity between two FC matrices as the correlation score in this manuscript. For the optimization, we use a genetic algorithm that iteratively combines multiple candidate input sets in order to form a near-optimal one (see Methods). To mitigate the lack of optimality guarantee, we run the algorithm multiple times. We denote the set of ROIs consistently selected across runs as the *consensus input set*.

We provide an illustration of the method based on simulated data in Figure 1C. First, we simulate 2000 time steps of a diffusion process driven by white noise on a graph composed of *n* = 10 nodes (*m* = 3 input nodes), with edge weights uniformly distributed between 0 and 1. Using these time series, we compute the associated FC matrix **F**_*emp*_. Then, we solve Problem 4 for *U* varying from 1 to *n*. We observe that the method retrieves the correct input set and produces an FC matrix that is similar to the empirical one.

### Linking the connectome to multiple functional states

We apply our approach and solve Problem 4 with empirical MRI data of 100 unrelated individuals. The data was acquired and preprocessed by the Human Connectome Project (HCP) consortium [29]. For each individual, we extract a connectome and FC matrices for resting-state and seven tasks (Figure 1B). The tasks are labelled as emotional processing, gambling, language processing, motor task, relational processing, social cognition and working memory. Each task required an active participation from the individuals. Although the properties of resting-state FC are known to be fundamentally different from that of task FC [33], we deliberately choose to treat resting-state in the same way as task conditions in order to test whether our approach is able to distinguish it. For simplicity, we refer to both resting-state and task conditions as *states* in the remainder of the manuscript. Finally, we use the brain parcellation introduced by Destrieux et al. [34] and composed of *n* = 164 ROIs including subcortical structures and cerebellum.

For the group-level analysis, we compute an average connectome and an average FC matrix **F**_*emp*_ for each state (see Methods). In order to study the stability of our results with respect to the number of input ROIs, we solve Problem 4 with *U* increasing from 1 to *n*. For each upper bound *U*, we define the consensus input set as the set of ROIs selected at least 25 times over 30 optimization runs.

Figure 2A shows the correlation score between **F**_*emp*_ and **F**_*SI*_ using the consensus input set. The curves increase with *U*, up to small drops due to the heuristic nature of the optimization (see Methods), until they reach a plateau at values ranging from *r* = 0.54 for resting-state to *r* = 0.7 for the motor task. We can compare these values with three baselines (see Methods for details about the baselines definition). The first one is the correlation score between **F**_*emp*_ and the adjacency matrix of the connectome. The second is the plateau correlation obtained by applying our approach to a randomly re-labelled connectome and keeping **F**_*emp*_ ordered, in order to break the ROI-to-ROI correspondence between structure and function while maintaining the network properties of the connectome. The third baseline is the maximum correlation score between **F**_*SI*_ and **F**_*emp*_ obtained with random input sets having the same average cardinality as the identified sets. In Figure 2A, we draw for each baseline the highest value across states and see that our approach produces a better matching for all states.

**Figure 2:**
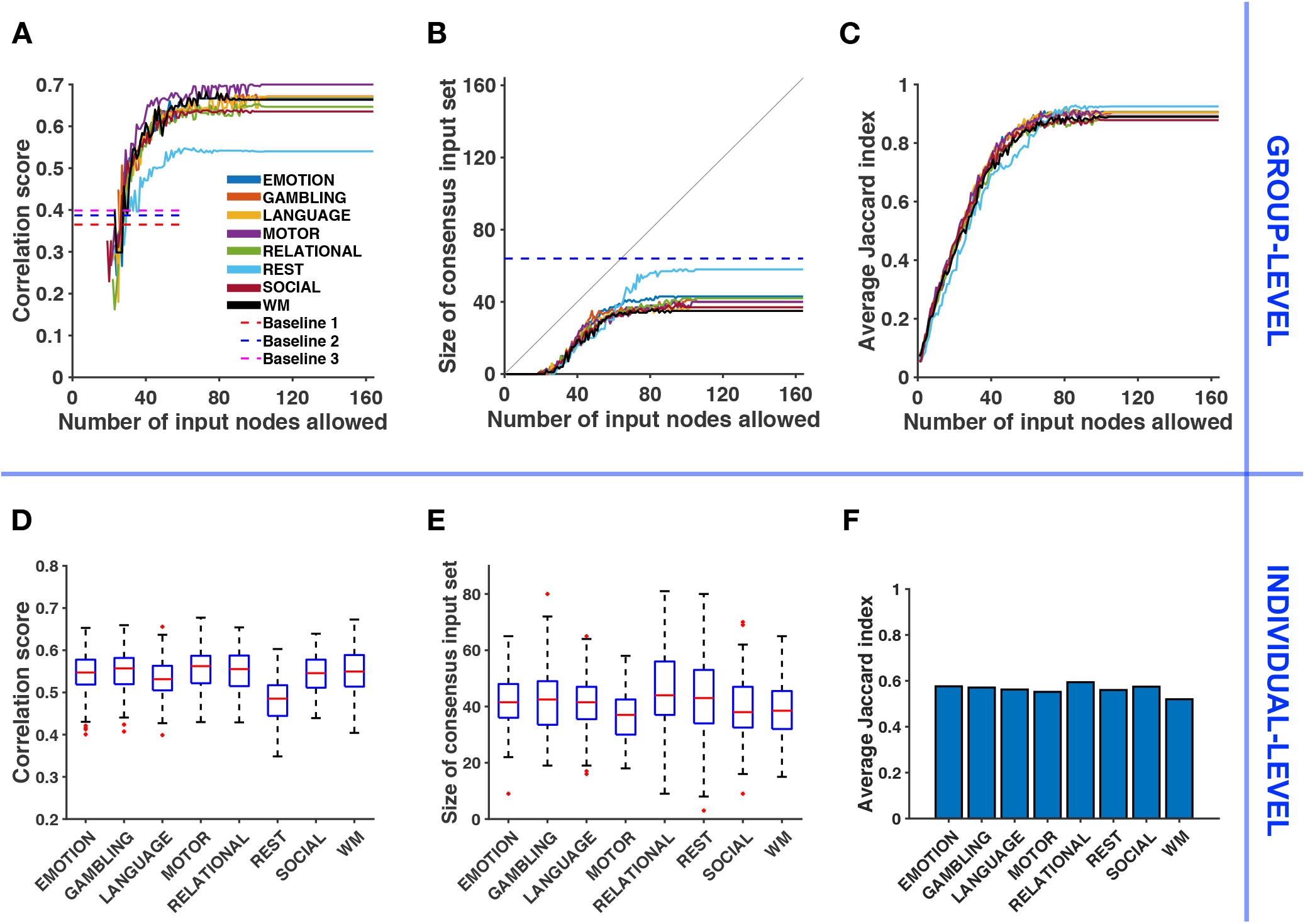
Relating structure-informed and empirical functional connectivity: Group-level analysis and individual-level variability. **A)** Correlation score between structure-informed and empirical functional connectivity with respect to the number of input nodes allowed *U* (group-level). **F**_*SI*_ is obtained using the consensus input set. Dashed lines represent baselines corresponding to the similarity between **F**_*emp*_ and (1) the adjacency matrix of the connectome, **F**_*SI*_ based on a re-labelled connectome and (3) **F**_*SI*_ obtained with a random input set (see Methods). **B)** Size of the consensus input set with respect to the number of input nodes allowed *U* (group-level). The grey line denotes the identity function *y* = *x*. The dashed blue line corresponds to the minimum number of input nodes selected for Baseline 2, over all conditions and all randomizations. **C)** Average Jaccard index between the 30 input sets identified by the optimization algorithm with respect to the number of input nodes allowed *U* (group-level). **D)** Variability across individuals of the correlation score between structure-informed and empirical correlation matrices, with *U* = *N*. **E)** Variability of the size of the corresponding consensus input set. **F)** Average Jaccard index of consensus input sets across individuals.

Figure 2B shows that the consensus input set is empty for all states until we allow the selection of at least 19 input nodes. Then its size stabilizes between *m* = 35 for the working-memory task (WM) and *m* = 58 for resting-state. These values are lower than the number of input nodes selected when applying our approach to a randomly re-labelled connectome (Baseline 2, *m* = 64, minimum across states and randomizations). To evaluate the consistency of identified input sets across optimization runs, we report in Figure 2C the evolution of the average Jaccard index. The Jaccard index *J* between two sets 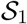 and 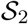 measures the overlap between these sets and is computed as

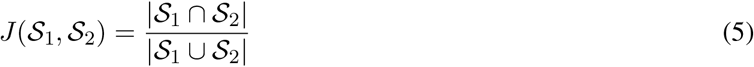

with *J* = 0 indicating no overlap and *J* = 1 indicating perfect overlap. The average Jaccard index 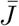 is computed over all pairs of the 30 optimised input sets. We observe that the method selects consistent input sets (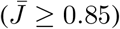) when *U* ≥ 70.

We also perform an individual-level analysis in the following way. We apply the method to each individual and set *U* = *N* in order to reduce the computational cost of the optimization. We therefore obtain one consensus input set and one correlation score for each individual and for each state. In Figure 2D, we notice that the correlation scores are lower than at the group-level, for all states. A repeated measures ANOVA determines that the mean correlation score differs significantly between states (*F* (7, 693) = 27.928, *p* < 10^−25^, Greenhouse-Geisser corrected). The variance in each condition does not significantly differ (Levene’s test, *p* > 0.5), and a post-hoc analysis after visual inspection confirms that the mean correlation score in resting-state is significantly different than in any task condition (Tukey’s HSD, *p* < 0.005). The post-hoc analysis also reveals that the mean correlation score significantly differs between the language task and the motor task (Tukey’s HSD, *p* < 0.005). Figure 2E shows the variability of the size of the consensus input set in the population. The relational processing task and the resting-state display a higher variability in the number of input ROIs selected than other states. In Figure 2F, we evaluate the variability of the consensus input set *in the population* by computing the average Jaccard index 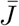 over all pairs of the 100 consensus input sets (one for each individual). We observe a moderate overlap of the consensus input set across individuals (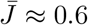, expected value of *J* for randomly chosen sets with cardinality *m* = 40: 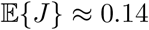, see Methods for the derivation).

### Analysis of input ROIs across functional subsystems

At the group-level, we turn our attention to the ROIs composing the input sets that we identified. In Figures 3A (motor task) and 3B (resting-state), we can follow the evolution of the number of selections of each ROI when the maximum cardinality *U* of the input set increases. We point out that when *U* is incremented, we perform the new optimization runs while ignoring previously computed solutions in order to assess the consistency of successively computed solutions. A first observation is that the selection of input ROIs is stable, that is once a region is selected it is typically selected again for higher values of *U*, as indicated by the horizontal red lines. Moreover, dark red pixels for a given ROI indicate that it is consistently selected across 30 independent optimization runs for a fixed *U*. We make a second observation by grouping ROIs according to the functional subsystems defined by Yeo et al. [35] and presented in Figure 3C (we include the cerebellum in the “subcortical” subsystem for visualization). Regions belonging to limbic and subcortical subsystems are selected together, up to some exceptions. These observations are also valid for the other tasks (corresponding figures are available in Supplementary material, Figure S1).

**Figure 3:**
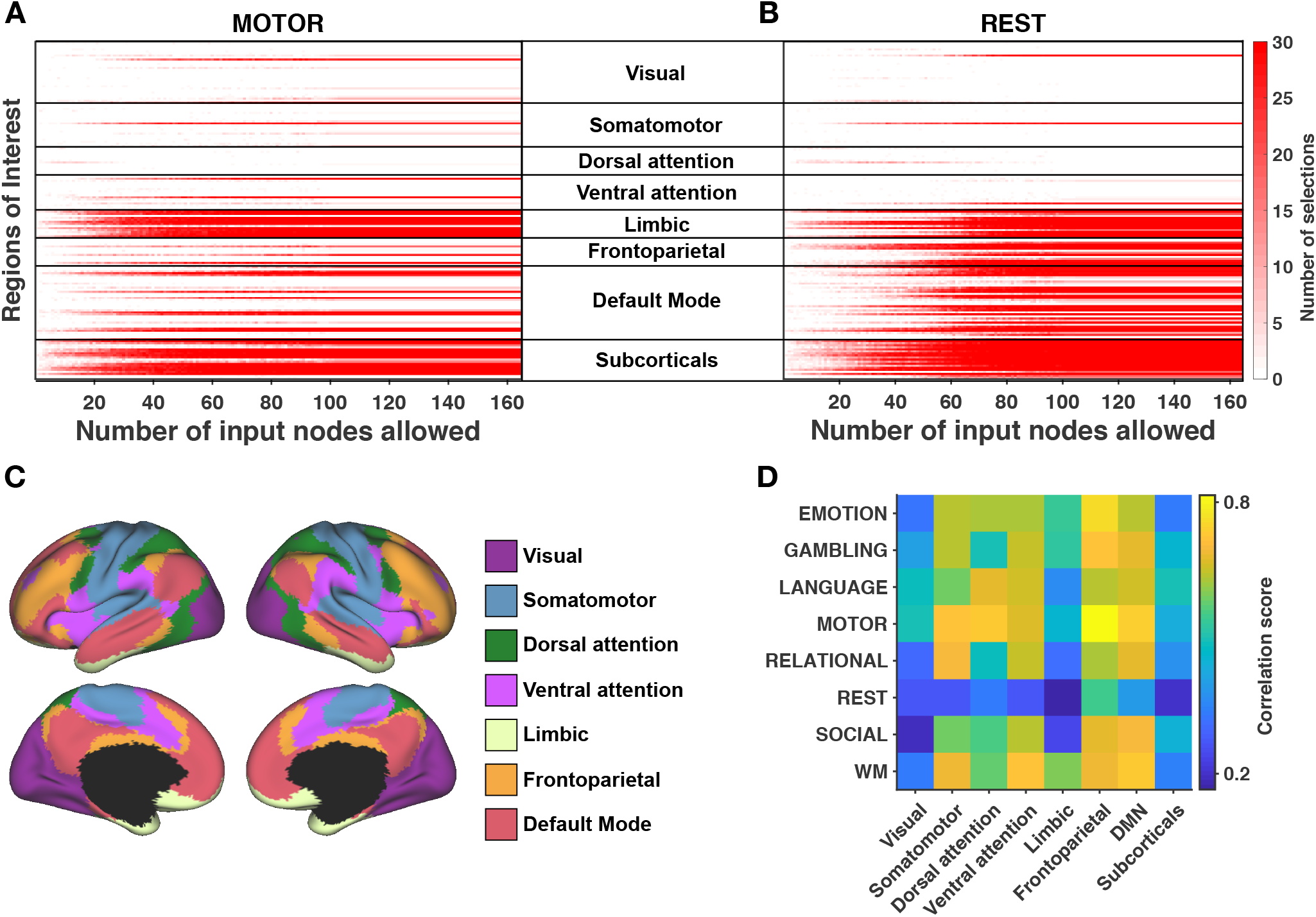
Analysis across functional subsystems (group-level). **A)** (resp. **B**) Evolution of the number of selections (from 0-white, to 30-red) of each Region of Interest (ROI) with respect to the number of input nodes allowed *U* for the motor task (resp. for resting-state). ROIs are arranged according to the functional subsystems described by Yeo and colleagues [35]. The cerebellum is included in the "subcorticals" subsystem for visualization and corresponds to the last two lines (left and right hemispheres). Corresponding figures for the other tasks are available in Supplementary Figure S1. **C)** Cortical localization of Yeo’s subsystems. **D)** Correlation between structure-informed and empirical functional connectivity with *U* = *N*, splitted into Yeo’s subsystems. Structure-informed functional connectivity is computed using the consensus input set.

Recent studies investigated how the connectome shapes functional connectivity at the level of subsystems and showed that the coupling between structure and function is stronger for some subsystems than others [36, 32, 37]. In Figure 3D, we use the consensus input sets identified at the group-level and compute the correlation score between the entries of **F**_*SI*_ and **F**_*emp*_ associated with the subsystems of Figure 3C. The results indicate that the association is the greatest in the frontoparietal lobe during the motor task (*r* = 0.82). Moreover, the visual and subcortical subsystems show low correlation scores for all states, while resting-state shows low correlation scores in all subsystems.

### Analysis of input ROIs across states

Next, we compare the composition of the identified input sets across states, at the group-level. Since we observed in Figure 2A that the correlation score reaches a plateau when *U* increases, we set *U* = *N* for this analysis. Moreover, we increase the number of optimization runs to 100 to evaluate more precisely the selection of each ROI. Thus, we obtain 100 input sets for each condition.

Figure 4A depicts the number of selections of each ROI across states. Blue lines indicate ROIs that have been selected at least 90 times for all states. These ROIs mostly correspond to subcorticals (accumbens nucleus, amygdala, hippocampus, pallidum, thalamus and subcallosal gyrus) and limbic regions (medial orbital sulcus, gyrus rectus and left suborbital sulcus). Regions of the default mode network (pericallosal sulcus, right suborbital sulcus and left posterior-ventral part of the cingulate gyrus) and of the somatomotor system (right paracentral lobule) complete the set of ROIs consistently selected across states. A cortical view of these regions is shown in Figure 4B. A table gathering the detailed numerical results by ROI is available at the end of the Supplementary materials and in the extended data.

**Figure 4:**
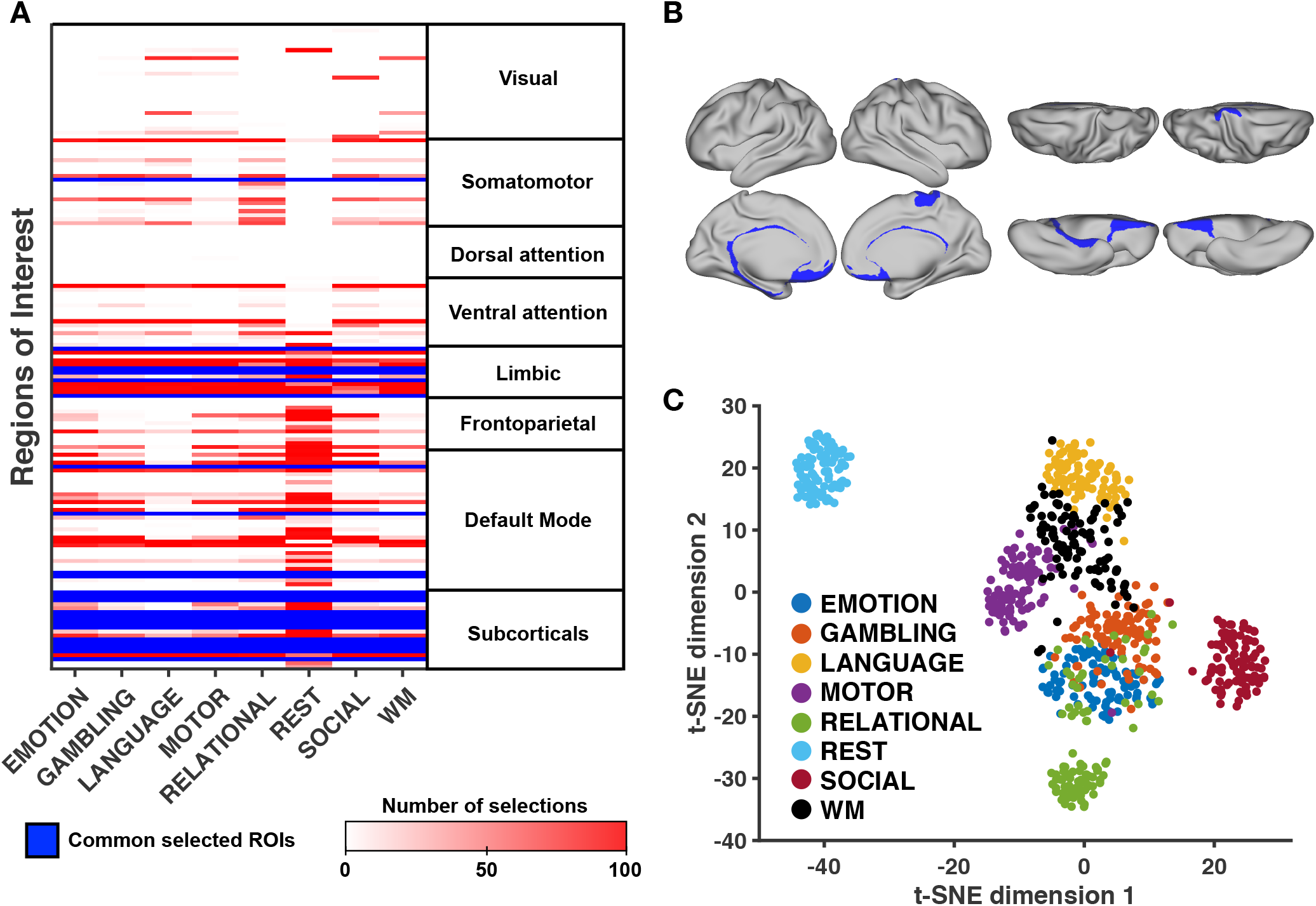
Analysis across functional states. **A)** Table summarizing the most frequently selected ROIs for each task. ROIs that are consistently selected at least 90 times over 100 runs for all functional states are highlighted in blue. ROIs are grouped according to Yeo’s functional subsystems. The cerebellum is included in the "subcorticals" subsystem for visualization and corresponds to the last two lines (left and right hemispheres) **B)** Cortical view of ROIs consistently selected across all tasks and resting-state. **C)** Two-dimensional projection of all input sets (100 runs, 8 states). We use the *t*-distributed Stochastic Neighbor Embedding algorithm (*t*-SNE, see Methods) in order to visualize the Jaccard similarity among all input sets. Each data point represents one such input set, and their proximity is proportional to their similarity.

In order to visualize the divergence of input sets across states, we use a dimensionality reduction method to project in two dimensions the *n*-dimensional binary vectors indicating which ROIs belong to each input set (*t*-distributed Stochastic Neighbor Embedding, see Methods). In Figure 4C, each data point represents one identified input set (100 runs, 8 states), and the proximity with each other is indicative of their overlap (Jaccard similarity). We distinguish clusters of points corresponding to different states. In particular, the cluster corresponding to resting-state is isolated. Among task conditions, there is a partial overlap of the clusters, with the input sets related to the social cognition task being more isolated from the others. A comparative cortical view of input ROIs for each condition is provided in Figure 5.

**Figure 5:**
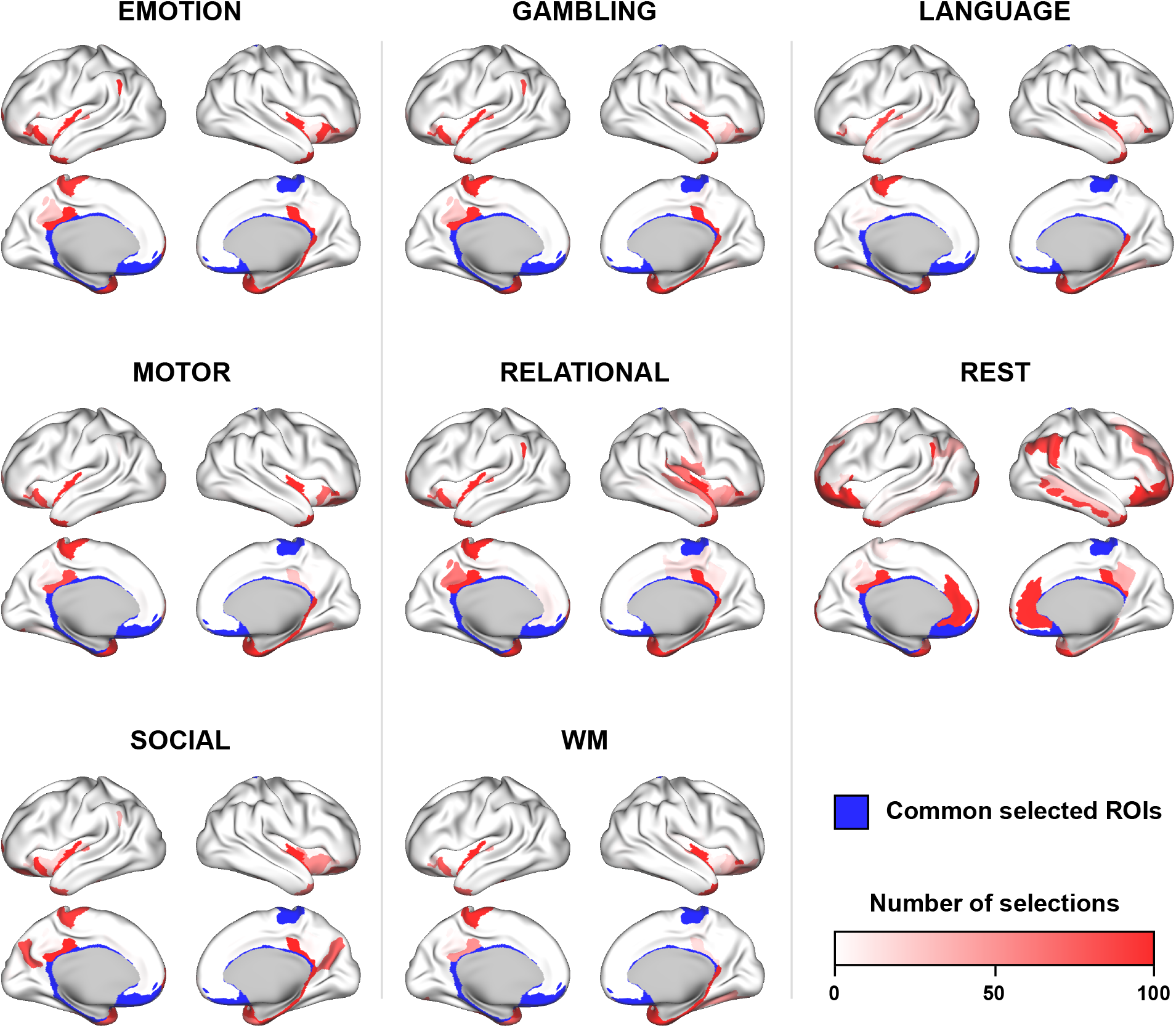
Cortical surface view of state-specific input ROIs. Over 100 runs of the optimization algorithm with *U* = *N*, we depict for each task the number of times each ROI is selected. Regions that were consistently selected across all states (≥ 90 selections) are shown in blue. Detailed numerical results by ROI are available at the end of the Supplementary materials, as well as in the extended data.

### Topological properties of input ROIs

In order to gain further insight into the topological properties of input ROIs in the connectome, we analyze the statistical association of the number of selections of each ROI with two nodal metrics: the weighted degree and modal controllability. The weighted degree of a node describes the strength of the connections with its neighbors, while modal controllability describes the ability of a node to drive the network towards hard-to-reach states requiring much control energy (see Methods and Refs [28, 18] for further details about modal controllability). For all tasks and resting-state, we report in Table 1 the correlation between these nodal metrics and the number of selections of ROIs. On the one hand, we find a significant inverse relationship between weighted degree and number of selections (lowest association: Spearman’s *ρ* = −0.3963 in resting-state, *p* < 10^−7^ for all states), which suggests that low-degree ROIs are selected more often. On the other hand, we find a significant proportional relationship between modal controllability and number of selections (lowest association: Spearman’s *ρ* = 0.4769 in resting-state, *p* < 10^−10^ for all states), indicating that ROIs having a high modal controllability are selected more frequently.

**Table 1:**
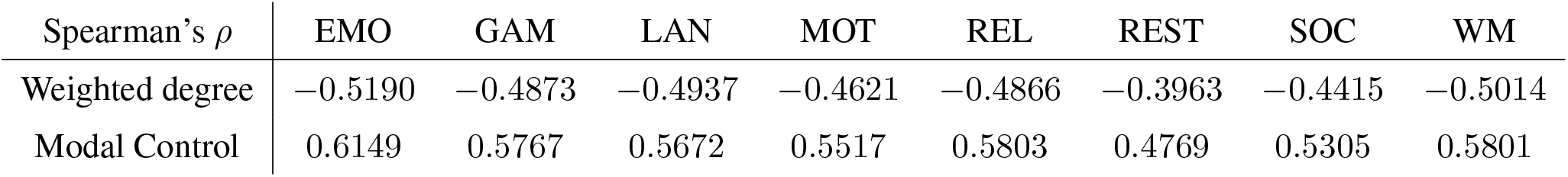
Spearman’s rank correlation between the number of selections of each ROI (out of 100 independent optimization runs, with *U* = *N*, group-level) and two nodal coefficients: the strength (weighted degree) and the modal controllability [28]. EMO: emotional processing, GAM: gambling, LAN: language processing, MOT: motor task, REL: relational processing, REST: resting-state, SOC: social cognition, WM: working-memory.

| Spearman’s $\rho$ | EMO | GAM | LAN | MOT | REL | REST | SOC | WM |
| --- | --- | --- | --- | --- | --- | --- | --- | --- |
| Weighted degree | -0.5190 | -0.4873 | -0.4937 | -0.4621 | -0.4866 | -0.3963 | -0.4415 | -0.5014 |
| Modal Control | 0.6149 | 0.5767 | 0.5672 | 0.5517 | 0.5803 | 0.4769 | 0.5305 | 0.5801 |

### Robustness of consensus input sets

Finally, we study the robustness of the link between structure-informed (**F**_*SI*_) and empirical (**F**_*emp*_) functional connectivity when the consensus input set is attacked. Here, an attack refers to the removal of a ROI from the initial input set, not from the connectome. We start from the correlations between **F**_*SI*_ and **F**_*emp*_ obtained at the group-level with *U* = *N* (Figure 2A). We progressively remove nodes from the consensus input set until it becomes empty. After each removal, we compute the correlation score obtained with the attacked input set. Since we previously observed that low-degree (resp. high modal controllability) ROIs are more likely to be part of the input set, the removal ordering is fixed by increasing order of weighted degree (resp. by decreasing order of modal controllability). In addition, we report the results related to 50 random removal sequences.

In Figure 6, we show the results for the motor task and the resting-state. We observe that the correlation score between **F**_*SI*_ and **F**_*emp*_ decreases slowly with the number of nodes removed from the consensus input set, no matter the removal ordering. For the motor task (resp. for resting-state), up to 75% (resp. 40%) of the nodes can be removed from the consensus input set before we reach correlation scores comparable to the three baselines previously defined (see Methods). Similar observations are valid for the other tasks (see Supplementary Figures S2 and S3).

**Figure 6:**
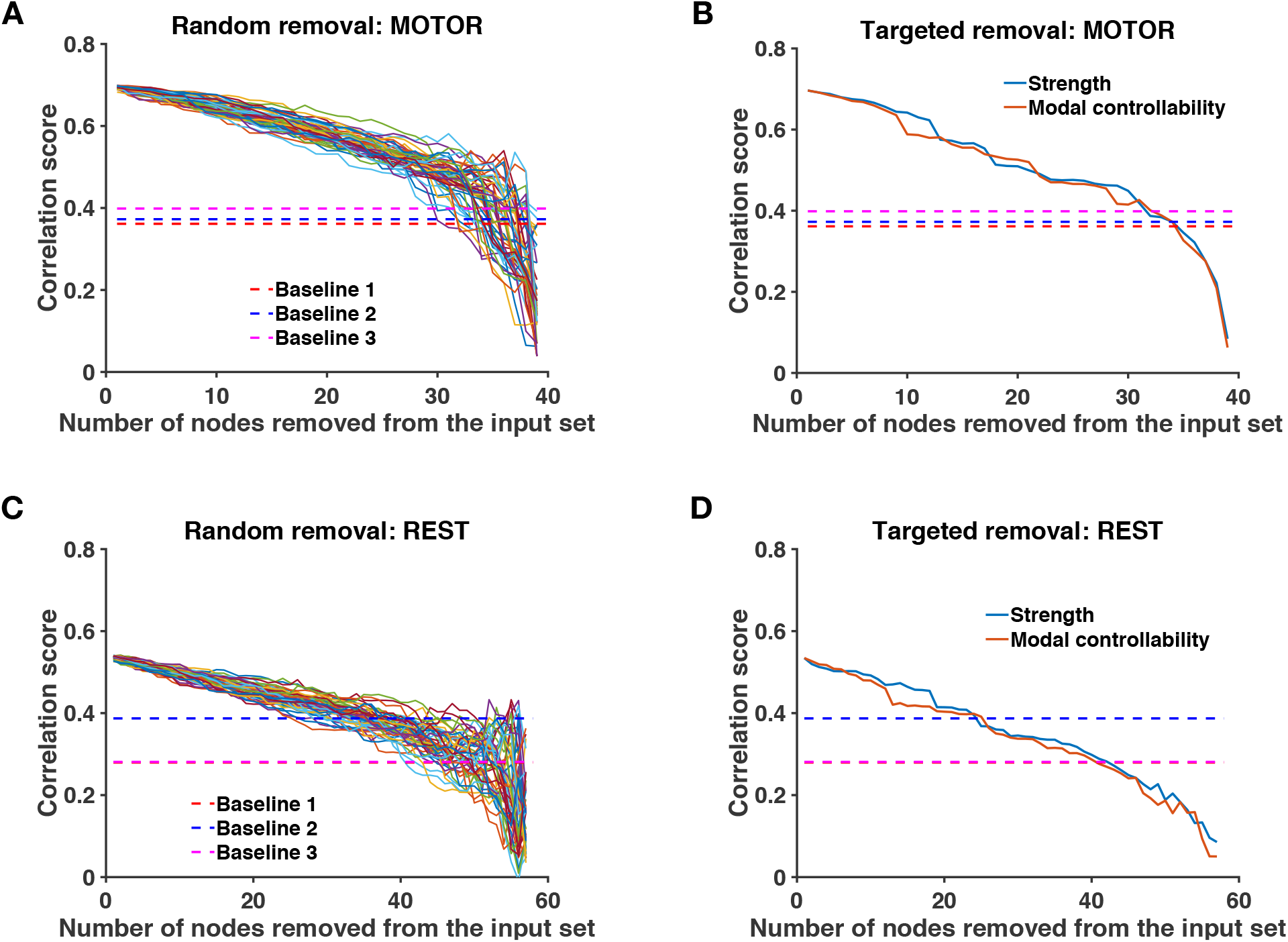
Robustness analysis. **A)**(resp. **C)**) Evolution of the correlation between structure-informed **F**_*SI*_ and empirical functional connectivity **F**_*emp*_ as a function of the number of ROIs removed from the consensus input set. Dashed lines represent the three baselines (see Methods), i.e. the correlation between **F**_*emp*_ and (i) the adjacency matrix of the connectome, (ii) **F**_*SI*_ based on a re-labelled connectome and (iii) **F**_*SI*_ obtained with a random input set. We consider 50 random removal orderings. **B)**(resp. **D)**) Same analysis, with removal ordering fixed by increasing weighted degree and decreasing modal controllability.

## Discussion

In this work, we studied the structure-function relationship in brain networks [8, 10, 11] across different task conditions as well as in resting-state. We showed that functional connectivity (FC), i.e. the coactivations among brain regions, can be explained by the correlations between the activities of these regions resulting from a linear dynamics spreading through the structure of the brain. This model, termed *structure-informed* FC, happens to be mathematically linked to the Gramian matrix used in controllability studies [28, 18, 30]. This provides a novel interpretation of FC in which we can leverage control theory to explain state-specific FC configurations arising from a fixed anatomical architecture. We thus proposed that different groups of regions controlling a diffusion dynamics through the wiring diagram of the brain are responsible for FC matrices corresponding to different states. We introduced a principled approach to test this hypothesis and found that sparse and stable groups of control regions, which partially overlap across states, generate FC matrices that are statistically comparable to empirical ones.

### Combining brain communication models and linear controllability

In this study, we considered a Laplacian diffusion dynamics to model the information flow between ROIs, following previous work [15]. While most studies on brain communication dynamics focused on the mechanisms governing the information flow in the absence of external intervention [16, 38, 39, 32, 40], it has been pointed out that the connectome is a controllable network capable of integrating context-dependent signals [20]. Earlier work on the controllability of the connectome typically fixed the transition matrix **A** (Equations 1-2) as the adjacency matrix of the connectome [28, 18, 30]. This is a common approach, but we should stress that it is possible to define multiple flow dynamics upon a fixed network structure [41]. Here, we suggest that combining a model of information flow [14] with the ability to modulate it through control regions [18] can improve our understanding of the structure-function relationship across functional states. Future research should investigate how other communication dynamics influence our results. For instance, we also tested our approach with **A** being the adjacency matrix of the connectome and obtained similar results with significant overlap in the computed consensus input sets (see Supplementary Table S1 for a comparison). Alternative dynamics such as decentralized (i.e. directed) brain communication models [16, 39] could provide complementary insights into the structure-function relationship in the human brain.

As in former connectomic studies [15, 28, 18, 30], our approach relies on linear time-invariant modelling (Equation 1). Despite the known non-linearities of neural dynamics [42], first-order approximations have been proved useful in capturing various aspects of brain functioning at different spatio-temporal scales [43, 44, 45]. In addition, time-invariance implies that the structure of the system does not evolve over time. Although the white matter architecture evolves over long timescales [46, 47], significant changes in the topology of the connectome are not expected over the duration of an MRI scan. Assuming linearity and time-invariance allowed us to derive an analytical expression of structure-informed FC (Equation 3). Since the heuristic optimization computes this matrix a large number of times in order to find a near-optimal input set, relying on an efficiently solved analytical expression of structure-informed FC rather than simulating the system at each iteration is computationally beneficial, although the computational cost remains a limitation of our framework. In sum, we argue that linear and time-invariant modelling of functional connectivity constitutes a reasonable and computationally tractable approach. Future studies are required to assess how much these assumptions can or should be relaxed in light of more realistic models compatible with the biology.

Similarly to previous studies in the field of brain communication dynamics and controllability, we considered in this work a coarse-grained parcellation spanning the entire brain [15, 18]. We suggest that the proposed method is also suitable at the level of subregions, provided that there exists an appropriate linear time-invariant model describing the dynamics at this scale, and we encourage future research in that direction.

### Well-defined sets of control regions drive state-specific functional connectivity

In our analyses, we identified sparse groups of regions that are thought to support the control of state-specific brain activity. The fact that our method finds such *sparse* input sets (Figure 2B), i.e. that empirical FC can be explained more simply from the true connectome structure than from a randomly re-labelled network, suggests the fitness of our model, in line with Occam’s razor principle. Our model also captured differences between states in terms of their respective input sets (Figure 4C), supporting the idea that different states are triggered by partially overlapping yet distinct sets of control regions. Because our approach involves a heuristic optimization algorithm, we assessed the consistency of the identification procedure (Figure 3A-B) and the robustness of the identified input sets (Figure 6). Moreover, we found that ROIs having low degree and high modal controllability, which are topological properties associated with the brain structure independently of any activation measure, have a higher probability to be part of an input set (Table 1). Together, these results suggest that the identified input ROIs play a central role in driving FC across the white matter wiring.

Importantly, this role does not imply that identified ROIs systematically match the active areas traditionally detected in fMRI analyses. For example, the primary motor cortex (M1) is not part of the input set of the motor task (Figure 5), although it displays strong activation in the functional data. This activation results from the fact that M1 forms a hub in the motor task, receiving projections from multiple regions, including the somatosensory and parietal cortices as well as premotor areas, and sending output commands to the periphery. This “centrality” however does not entail that M1 is part of the set of drivers that put the brain in a state that is suitable for motor control. In this regard, our findings are supported by recent experimental evidence in mice showing that thalamic inputs are essential to drive the motor cortex during movement execution [48]. A similar example is that of Wernicke’s area, which was not part of the input set of the language processing task (Figure 5) but whose activation is often associated with language understanding. More generally, the fact that drivers are preferentially ROIs with low degree and high modal controllability is consistent with the idea that reaching demanding states requires the control of decentralized and distributed areas, which in turn influence the whole system including hubs, such as M1 or Wernicke’s area [21, 18, 26, 27, 49].

In order to gain a better insight into the role of the ROIs that we identified, we turn our attention to the drivers common to all states. The presence of subcortical structures (including basal ganglia, amygdala, hippocampus and thalamus) in the input set of all states is consistent with their strong contribution to whole-brain communication [50], motor control [51], language processing [52], reward-related processing [53] and cognition in general [54]. Anatomical and physiological evidence established the existence of cortico-subcortical loops supporting functionally segregated systems [55]. Within these loops, which include the anterior cingulate and dorsolateral prefrontal cortices that have been designated as cognitive control centers [24, 56, 25], subcortical structures are thought to modulate the process of action selection, given afferent cortical signals [54]. Regarding the other identified regions, we provided in our analyses a numerical assessment of their consistency in the context of our model. Their functional relevance remains to be further validated in neurophysiological studies involving tailored experimental protocols, and the present work can guide future research investigating brain regions that underlie task-specific control.

### Distinguishing resting-state from task conditions

In this study, we applied our approach to both resting-state and task-based FC without *a priori* distinction, although their properties are different [33] and resting-state was the only condition that did not require any active involvement of the individuals. Interestingly, our method captured the singularity of resting-state in several regards: the matching between structure-informed and empirical FC is lower (Figure 2A-D) and requires more input regions (Figure 2B-E). Moreover, the input set related to resting-state is distinct from that of task conditions (Figure 4C) and includes more regions belonging to the frontoparietal subsystem and to the default mode network (Figure 4A).

Accumulating evidence from fMRI studies speculate that resting-state FC forms a “standard” architecture in which segregated functional subsystems are represented, and which supports the transfer of information related to the implementation of tasks [57, 58, 35, 33 7, 59]. This could explain why, from a controllability viewpoint, our results distinguish rest (the passive, default state) from task conditions (the active, target states). Following the hypothesis that resting-state connectivity supports task implementation, an extension of this study consists in applying our framework to the graph structure defined by resting-state FC instead of the connectome, in order to investigate which brain regions drive the rest-to-task transitions.

### Conclusion and future work

This report presented a system-theoretic framework for identifying potential state-specific control regions through a model linking structure and function in human brain networks. In this respect, it linked concepts of brain communication dynamics and connectome controllability. We expect that future research, for instance in clinical populations, will further validate the proposed approach by studying the impact of neurological deficits and lesions on the identified control regions. This work could in turn guide physiological studies investigating the role of particular regions in controlling brain processes. Future work should also analyze individual differences in the identified control regions and their possible relation to behavior.

## Methods

### Dataset

We retrieved the preprocessed “100 unrelated subjects” dataset of the Human Connectome Project (HCP) database (https://db.humanconnectome.org/), HCP 1200 release [29]. All individuals (54 females, 46 males, 22-36 y.o.) gave written informed consent to the HCP consortium. Scanning protocols were approved by the local Institutional Review Board at Washington University in Saint Louis. Acquisition parameters are detailed in previous HCP reports [60, 29, 61]. Preprocessing consisted of HCP minimal preprocessing pipelines [61]. We applied further processing steps in agreement with previously published studies using HCP data [49, 32, 62].

#### Parcellation

We used the cortical parcellation introduced by Destrieux and colleagues [34] and composed of 148 non-overlapping Regions of Interest (ROIs). Subcortical structures (thalamus, caudate nucleus, putamen, pallidum, hippocampus, amygdala, accumbens nucleus) and cerebellum were extracted using the FMRIB Software Library [63] and added to the parcellation for completeness, bringing the number of ROIs to *n* = 164.

#### Connectome reconstruction

The processing of diffusion data was conducted for each individual using state-of-the-art methods implemented in the MRtrix3 toolbox [64]. In summary, a tissue-segmented image was generated (MRtrix command `5ttgen`) in order to perform Anatomically-Constrained Tractography [65]. Then, multi-shell, multi-tissue response functions were computed (MRtrix command `dwi2response msmt_5tt`) in order to inform the Constrained Spherical Deconvolution (MRtrix command `dwi2fod msmt_csd`) [66]. Probabilistic tractography (MRtrix command `tckgen`) was performed using a second-order integration over fiber orientation distributions (iFOD2 method [67]) to allow for a more precise fiber tracking through crossing regions. This produced an initial tractogram composed of 10 millions streamlines that was corrected (SIFT2 approach [68], MRtrix command `tcksift2`) in order to obtain a more biologically meaningful representation of white matter tracts by computing an appropriate cross-sectional area multiplier for each streamline. Eventually, we built the adjacency matrix **S** of the connectome by computing the fiber density between each pair of previously defined ROIs (MRtrix command `tck2connectome` with option `-scale_invnodevol`). The group-average adjacency matrix is obtained as the entrywise average of the *K* = 100 individual-level adjacency matrices. Both group-average and individual matrices were kept unthresholded.

#### Empirical functional connectivity

We included fMRI data acquired during resting-state and seven tasks: emotional processing (EMOTION), gambling, language processing (LANGUAGE), motor response (MOTOR), relational processing (RELATIONAL), social cognition (SOCIAL) and working memory (WM) [29]. Resting-state Blood-Oxygenation-Level Dependent (BOLD) time series were filtered in forward and reverse directions (1st-order Butterworth, bandpass = [0.001,0.08] Hz) [69]. We did not regressed out the global signal. For both resting-state and task fMRI, the voxel time series were then z-scored and averaged in each ROI using the Connectome Workbench toolbox [70] and excluding outlier time points outside 3 standard deviations from the mean (Workbench command `-cifti-parcellate`). Empirical functional connectivity (FC) matrices **F**_*emp*_ were obtained by computing Pearson’s correlation coefficient between each pair of resulting time series. For each task, FC matrices of both fMRI phase encoding directions (left-to-right and right-to-left) were averaged in order to reduce the effect of artifactual noise. For resting-state, the four resulting matrices (2 scans, 2 phase encoding directions) were averaged for the same reason. The group-average FC matrix (for each task and resting-state) is obtained as the entrywise average of the *K* = 100 individual-level FC matrices. Both group-average and individual matrices were kept unthresholded.

### State correlation matrix of a linear diffusion process driven by white noise

We consider the following linear discrete-time invariant dynamics:

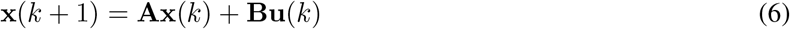

We excite the system with white noise signals **u**, and we assume that **x** is centered, **A** is stable, the input signals are not correlated with the initial state of the system (i.e. 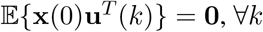) and input signals in **u** have unit variance. We compute the steady-state covariance matrix **Σ**= Cov(**x**) of System (6) as follows:

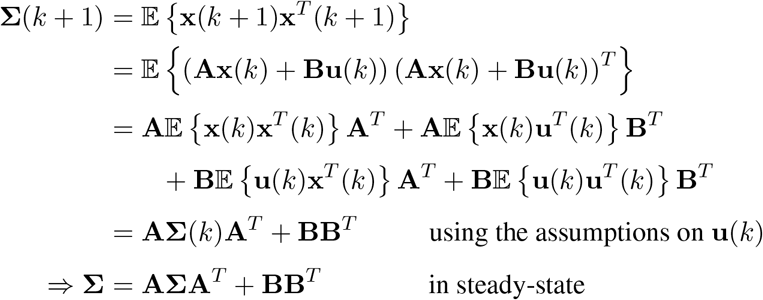

We notice that excitation signals with non-unit variance would result in a scaling of matrix **B**, which would not affect further results.

Defining **P** as the diagonal matrix containing only the diagonal entries of **Σ**(i.e. the states variances), we can apply a symmetric normalization to the steady-state covariance matrix to obtain a pairwise correlation matrix that we use as a model of functional connectivity:

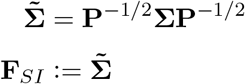

In this study, we consider the particular case of a diffusion process unfolding on the connectome, as proposed by Abdelnour and colleagues [15]. The state transition matrix **A** has therefore the following form:

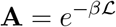

Here, with **D** being the diagonal matrix of the weighted degree of the ROIs, the matrix 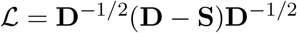 is the normalized Laplacian of the connectome. The parameter 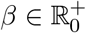 accounts for the sampling time with respect to the associated continuous-time system. While its value can be chosen arbitrarily, we choose here to fix it a priori to *β* = 0.72s in order to match the sampling rate of the empirical fMRI data used in this work [29]. In order to stabilize System (6), as done in several studies [18, 71] (see the detailed discussion in [30]), the entries of matrix **A** were further divided by 1 + *λ*_max_(**A**), where *λ*_max_(**A**) is the largest eigenvalue of **A**. In our case, the smallest eigenvalue of 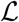 is always 0 since the Laplacian of an unsigned graph is positive semi-definite, and always possesses a zero eigenvalue [72]. Therefore, 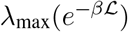 is always equal to 1.

### Finding the optimal set of control regions: Genetic Algorithms

Given the combinatorial nature of the optimization Problem 4, we must resort to heuristic methods in order to approach optimal solutions, without guarantee of optimality. A convenient choice is the family of genetic algorithms [73]. Here, the steps involved in the genetic algorithm that we used are *i*) generating a random population of admissible input sets, *ii*) selecting the best input sets in the population, *iii*) breeding a new generation of solutions by crossovers between selected input sets, *iv*) applying random modifications in the new population to avoid getting trapped in a local optimum and *v*) repeating the process until no more improvement is achieved after a given number of iterations. In the present study, we used the Matlab implementation of genetic algorithms, from the Global Optimization Toolbox, with default options and parameters. The Matlab code used to produce the results in this report is available online (https://github.com/bchiem42/Structure-informed-FC).

### Baselines

In order to assess how well our approach maps structure to function, we provide three baseline values.

#### Baseline 1

This is the Pearson’s correlation coefficient between the vectorized upper-triangular of the adjacency matrix of the connectome **S** and the empirical FC matrix **F**_*emp*_, without any transformation.

#### Baseline 2

We randomly re-label the ROIs of the connectome matrix **S** while keeping **F**_*emp*_ unchanged, and then apply our method. This null-model breaks the ROI-to-ROI correspondence between structure and function and preserves all network properties of the connectome. In the results, we report the maximum correlation score obtained over 30 random re-labelling.

#### Baseline 3

In order to assess the usefulness of solving the optimization Problem 4 to identify optimal input sets, we compute the correlation score between **F**_*emp*_ and **F**_*SI*_ obtained with an input set drawn uniformly at random, with cardinality *m* ∈ [37, 43], following the result depicted in Figure 2B. In the results, we report the maximum correlation score obtained over 30 random input sets.

### Modal controllability

Given a system defined on a network of *n* nodes, modal controllability is a nodal property that quantifies the ability of a single node to steer the system towards states requiring substantial input energy [28, 18, 30]. We compute the modal controllability *ϕ*_*i*_ of node *i* from the eigenvalues *λ* and the eigenvectors **v** of the adjacency matrix **S** of the connectome:

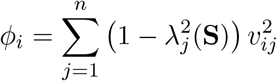

In this work, we computed modal controllability using the Matlab implementation provided by the authors of [18].

### 2D visualization of input sets using *t*-distributed Stochastic Neighbor Embedding

In a network of *n* nodes, we represent an input set as an *n*-dimensional binary vector indicating which node is selected (1) or not (0). In order to visualise how multiple input sets relate to each order, we can use dimensionality reduction to embed the *n*-dimensional vectors in two dimensions. In particular, the *t*-distributed Stochastic Neighbor Embedding [74] aims at finding a low-dimensional representation of high-dimensional vectors while preserving their local structure, such that similar vectors are represented by close points in 2D and vice-versa, with high probability. In this work, we used the Jaccard index to measure the similarity between vectors.

### Expected value of Jaccard index

We consider a set of *n* elements from which we draw two subsets 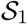 and 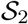 having the same cardinality *m* and whose elements are chosen uniformly at random. We denote the number of common elements between 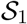 and 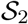 as 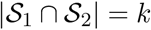. The corresponding Jaccard index is

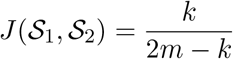

Now, the probability that the number of common elements is exactly *k* is

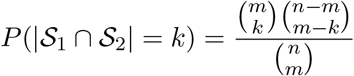

since we have 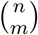 choices for the elements of 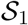 and 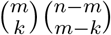 choices left for the elements of 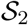. Therefore, the expected value of Jaccard index between two random sets of size *m* drawn from *n* elements is

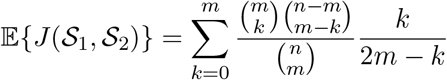

For *n* = 164 and *m* = 40 (see Figure 2E), we obtain 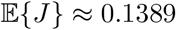.

## Acknowledgments

The authors would like to thank Laurence Dricot for her helpful comments and suggestions. Benjamin Chiêm is a FRIA (F.R.S.-FNRS) fellow. Data were provided by the Human Connectome Project, WU-Minn Consortium (Principal Investigators: David Van Essen and Kamil Ugurbil; 1U54MH091657) funded by the 16 NIH Institutes and Centers that support the NIH Blueprint for Neuroscience Research; and by the McDonnell Center for Systems Neuroscience at Washington University.

## Supplementary materials

**Table S1:** Comparison between two linear dynamics: the Laplacian diffusion dynamics used in the main manuscript (DIFF), and the dynamics defined by fixing the transition matrix **A** to the adjacency matrix of the connectome (ADJ). In both cases, we performed 100 runs of the optimization algorithm, with *U* = *N*. The Jaccard index *J* is computed between the consensus input sets (>= 90 selections) obtained with the two dynamics.

|  | EMO | GAM | LAN | MOT | REL | REST | SOC | WM |
| --- | --- | --- | --- | --- | --- | --- | --- | --- |
| Mean correlation score : DIFF | 0.66 | 0.67 | 0.68 | 0.7 | 0.65 | 0.54 | 0.63 | 0.68 |
| Mean correlation score : ADJ | 0.7 | 0.7 | 0.69 | 0.71 | 0.7 | 0.62 | 0.68 | 0.69 |
| $J(\text{DIFF}, \text{ADJ})$ | 0.92 | 0.88 | 0.88 | 0.87 | 0.92 | 0.86 | 0.77 | 0.93 |

**Figure S1:**
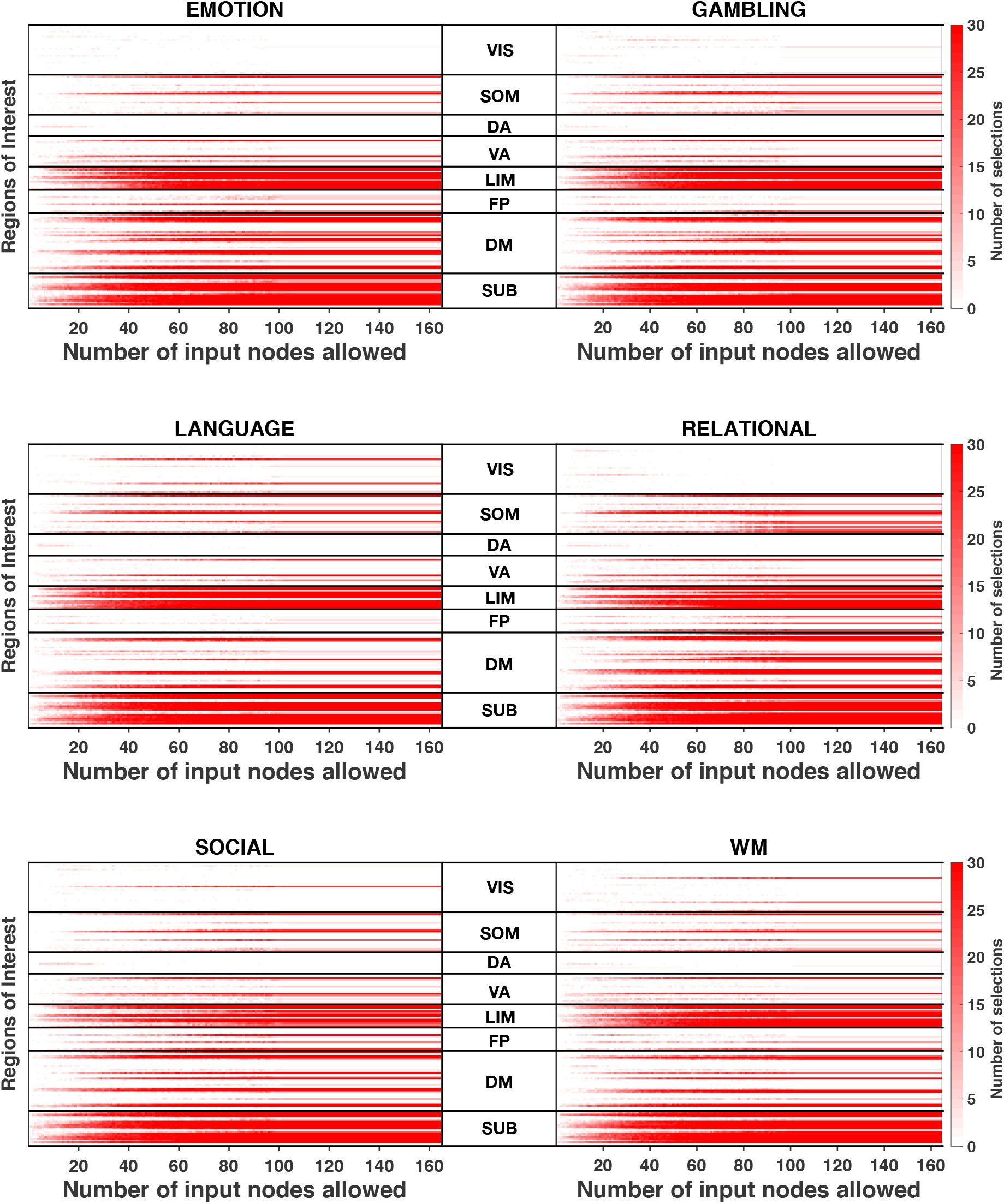
Analysis across functional subsystems (group-level). Evolution of the number of selections (from 0-white, to 30-red) of each Region of Interest (ROI) with respect to the number of input nodes allowed *U* for different tasks (MOTOR and REST are presented in the main manuscript). ROIs are arranged according to the functional subsystems described by Yeo and colleagues [35]. The cerebellum is included in the "subcorticals" subsystem for visualisation and corresponds to the last two lines (left and right hemispheres)

**Figure S2:**
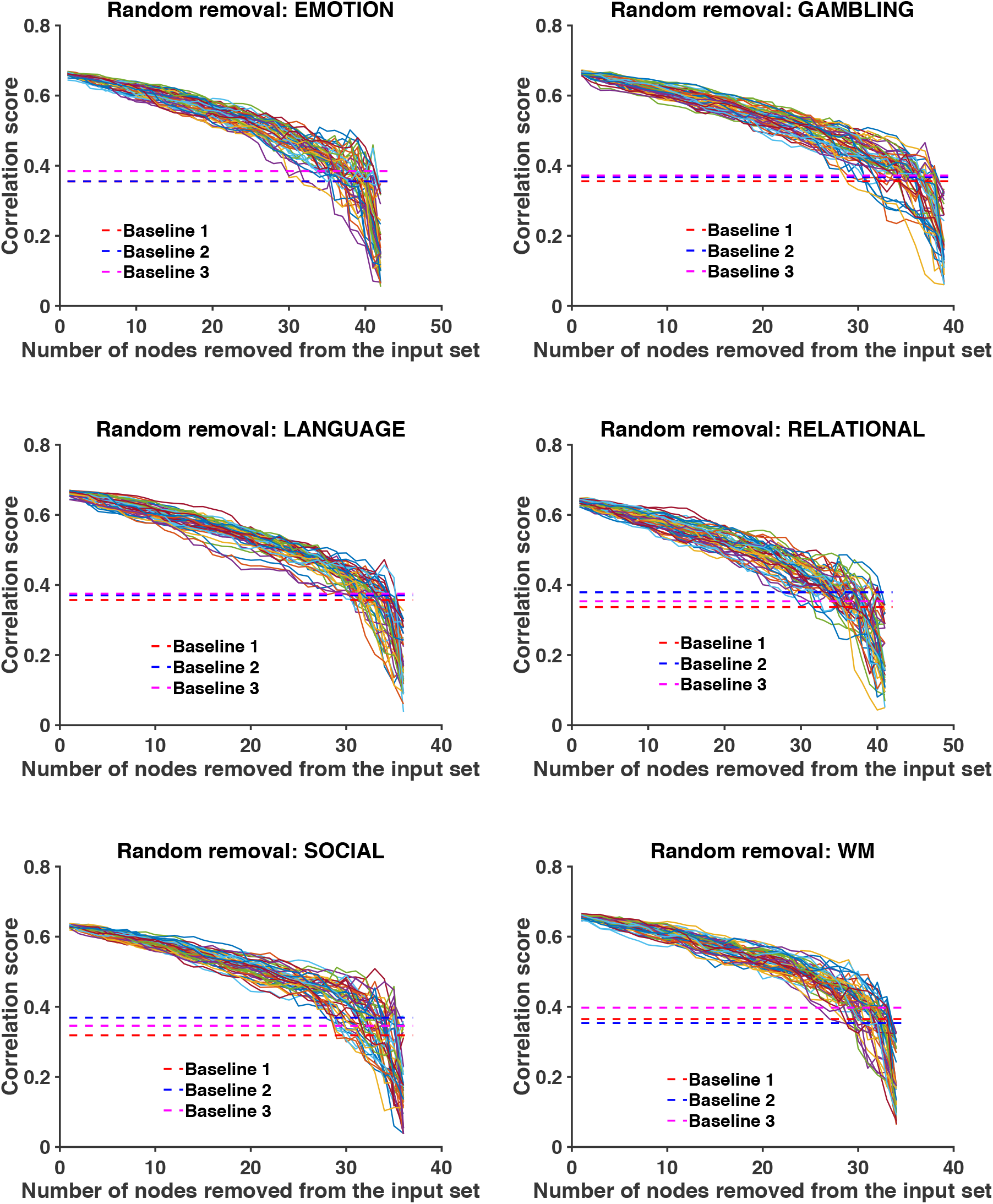
Robustness analysis. Evolution of the correlation between structure-informed **F**_*SI*_ and empirical functional connectivity **F**_*emp*_ as a function of the number of ROIs removed from the consensus input set. Dashed lines represent the three baselines, i.e. the correlation between **F**_*emp*_ and (i) the adjacency matrix of the connectome, (ii) **F**_*SI*_ based on a re-labelled connectome and (iii) **F**_*SI*_ obtained with a random input set. We consider 50 random removal orderings.

**Figure S3:**
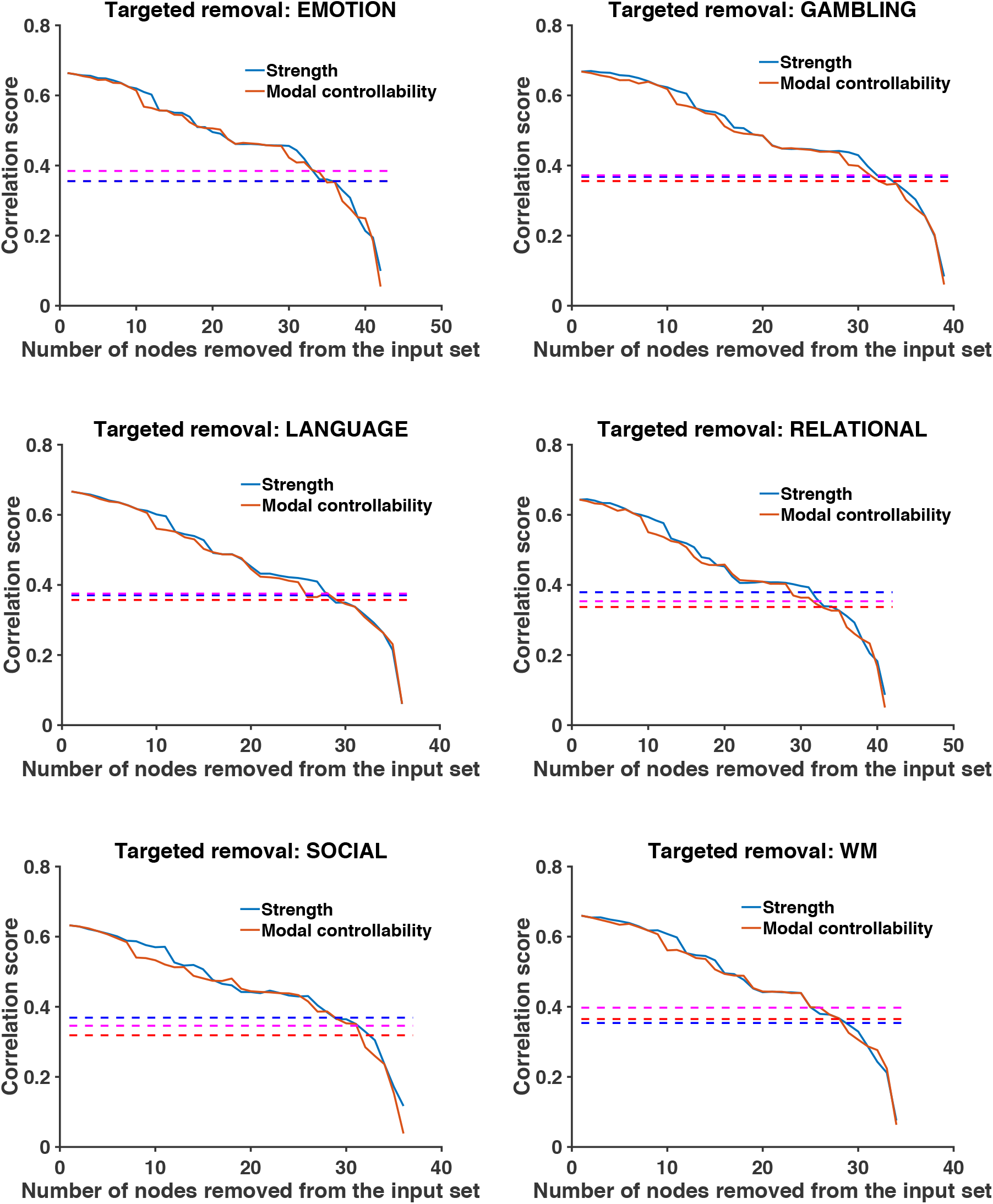
Robustness analysis. Evolution of the correlation between structure-informed **F**_*SI*_ and empirical functional connectivity **F**_*emp*_ as a function of the number of ROIs removed from the consensus input set. Dashed lines represent the three baselines, i.e. the correlation between **F**_*emp*_ and (i) the adjacency matrix of the connectome, (ii) **F**_*SI*_ based on a re-labelled connectome and (iii) **F**_*SI*_ obtained with a random input set. The removal ordering is fixed either by increasing weighted degree or decreasing model controllability.

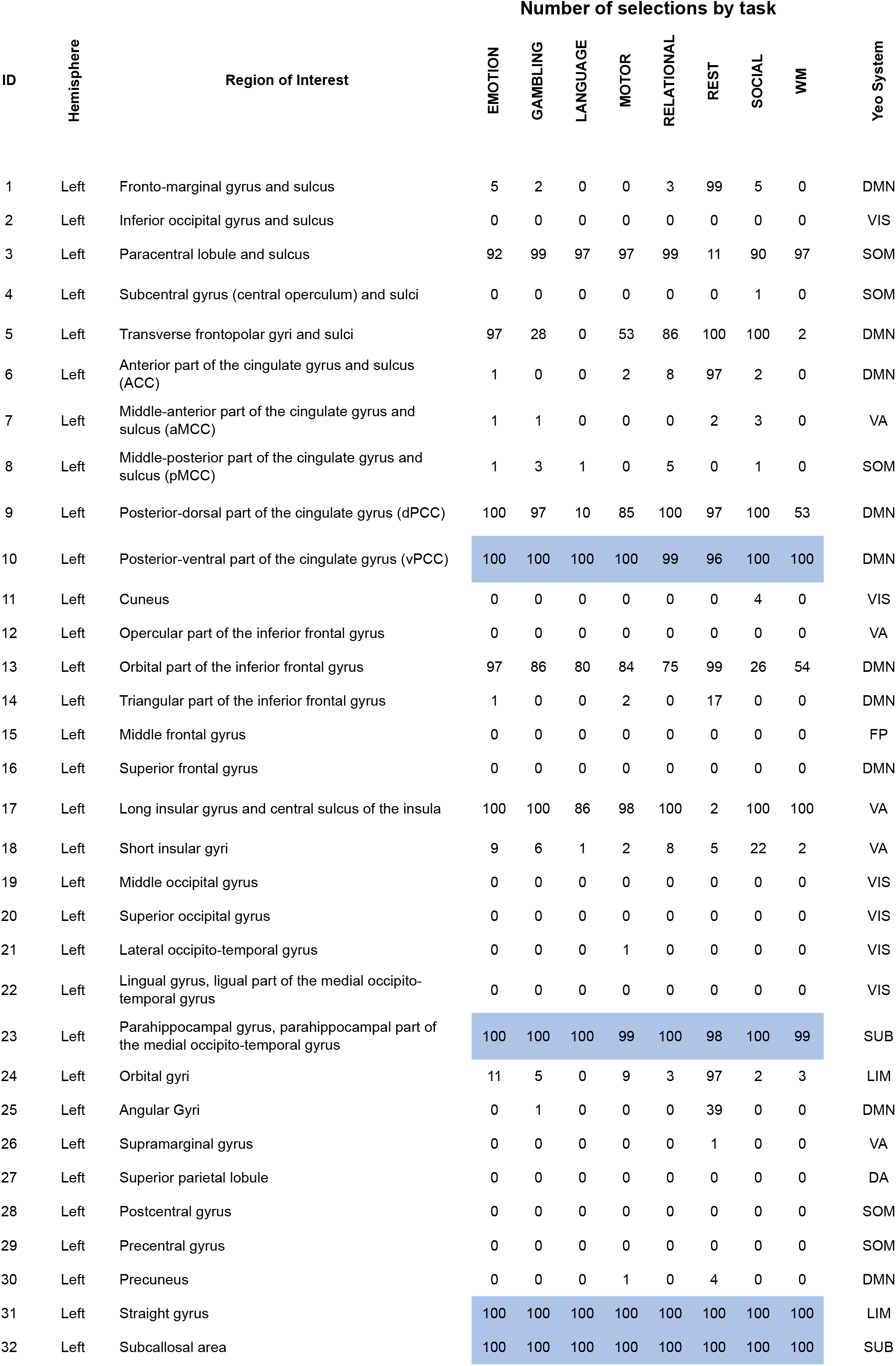
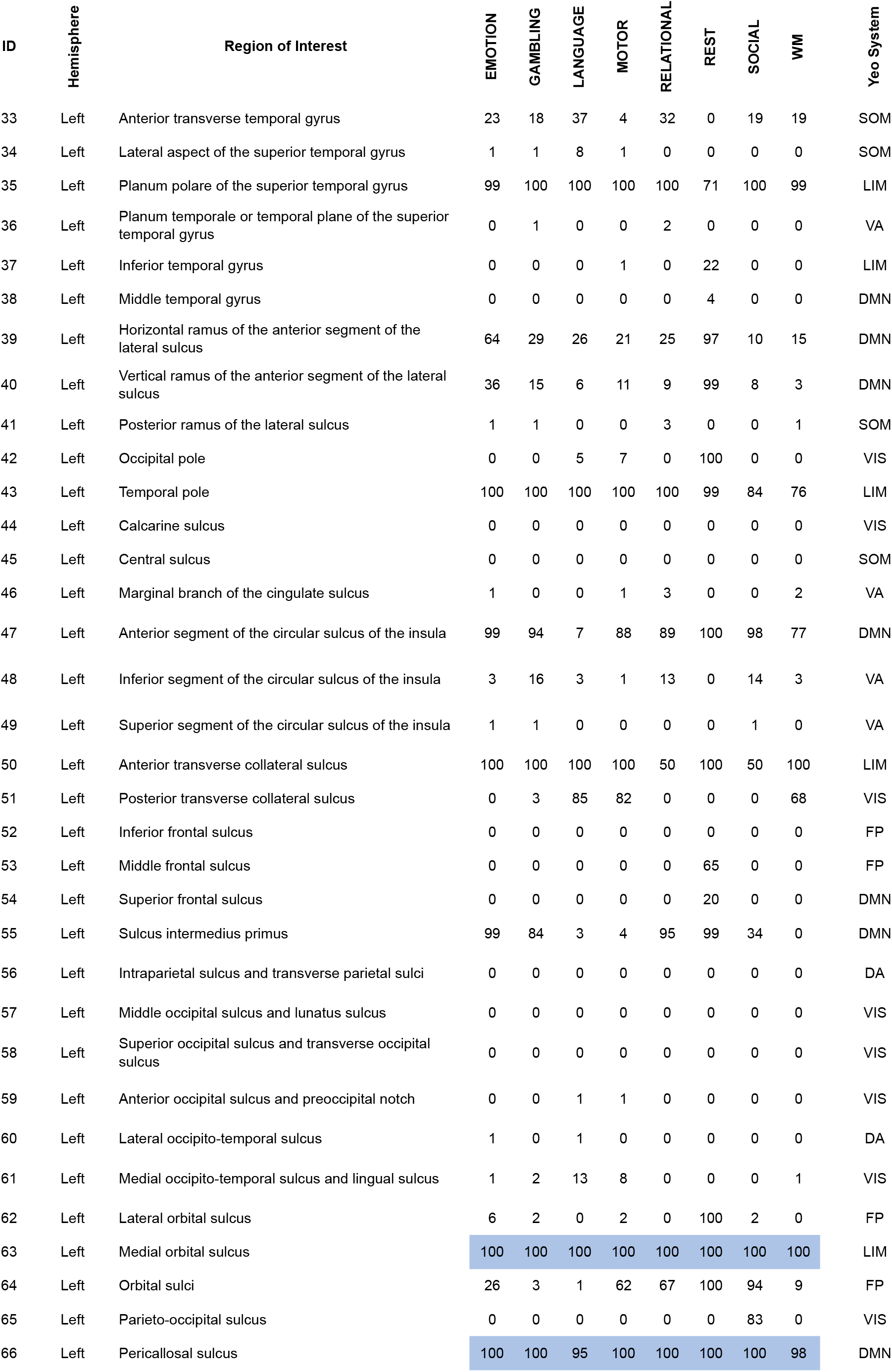
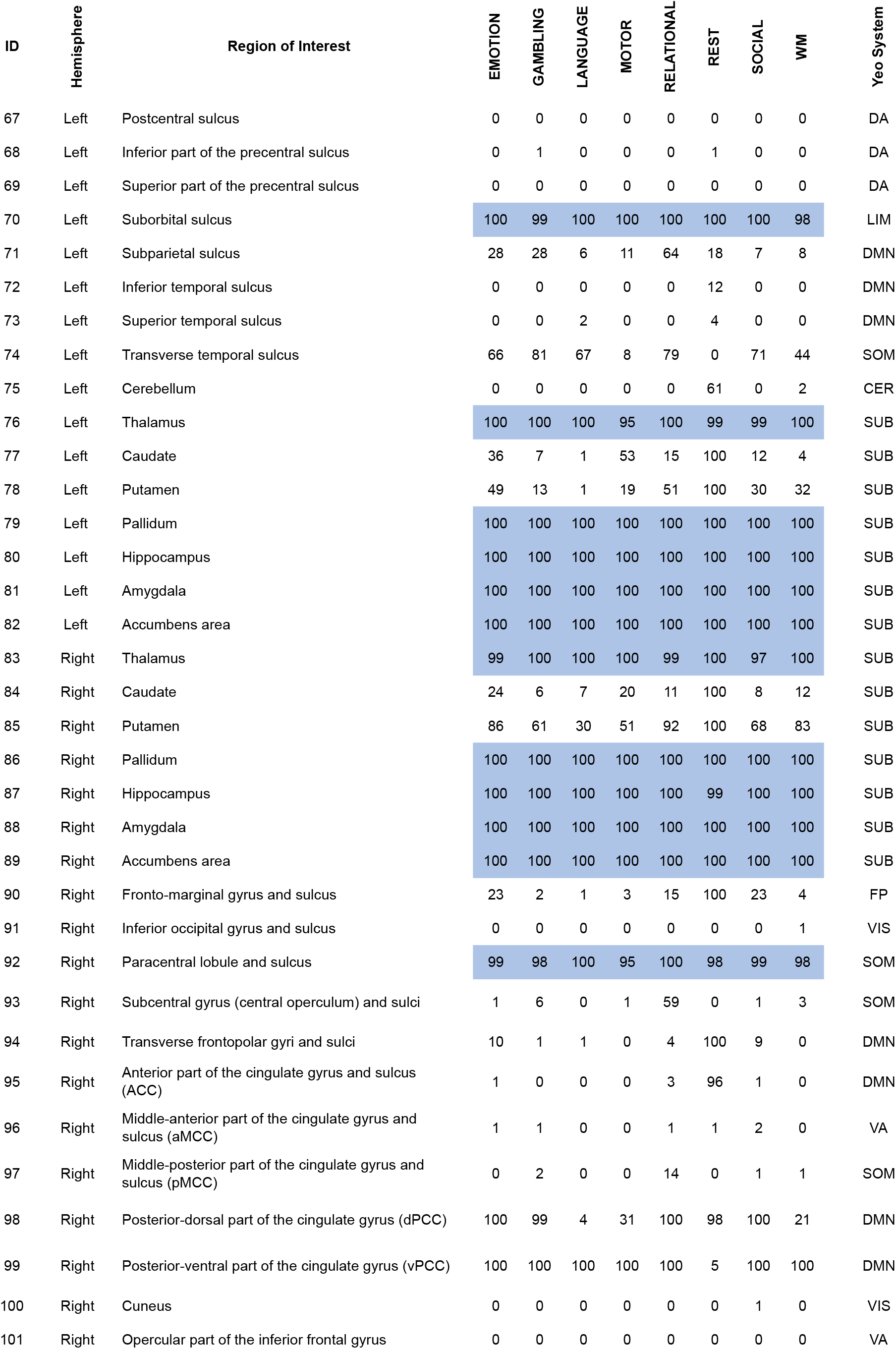
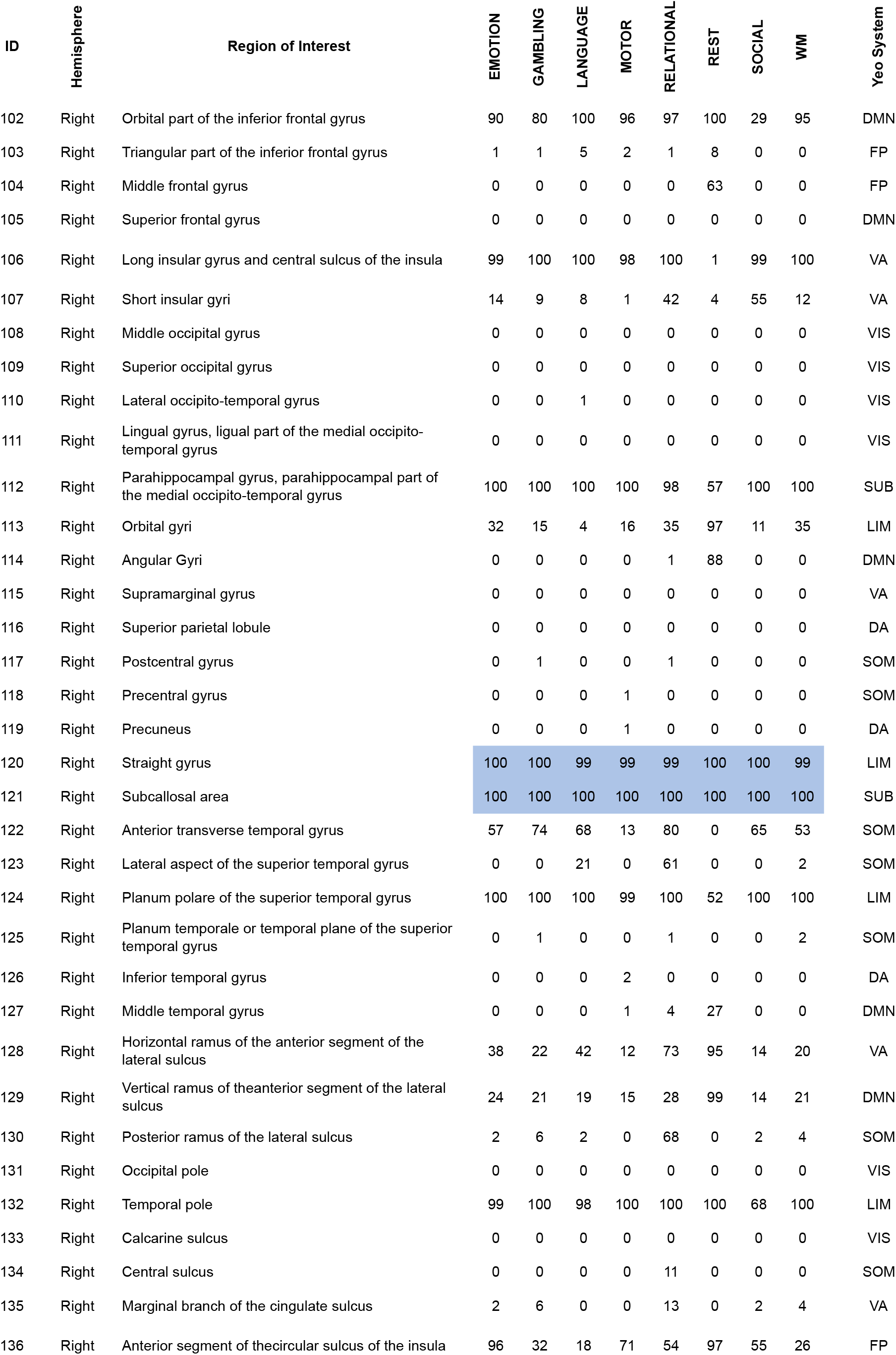
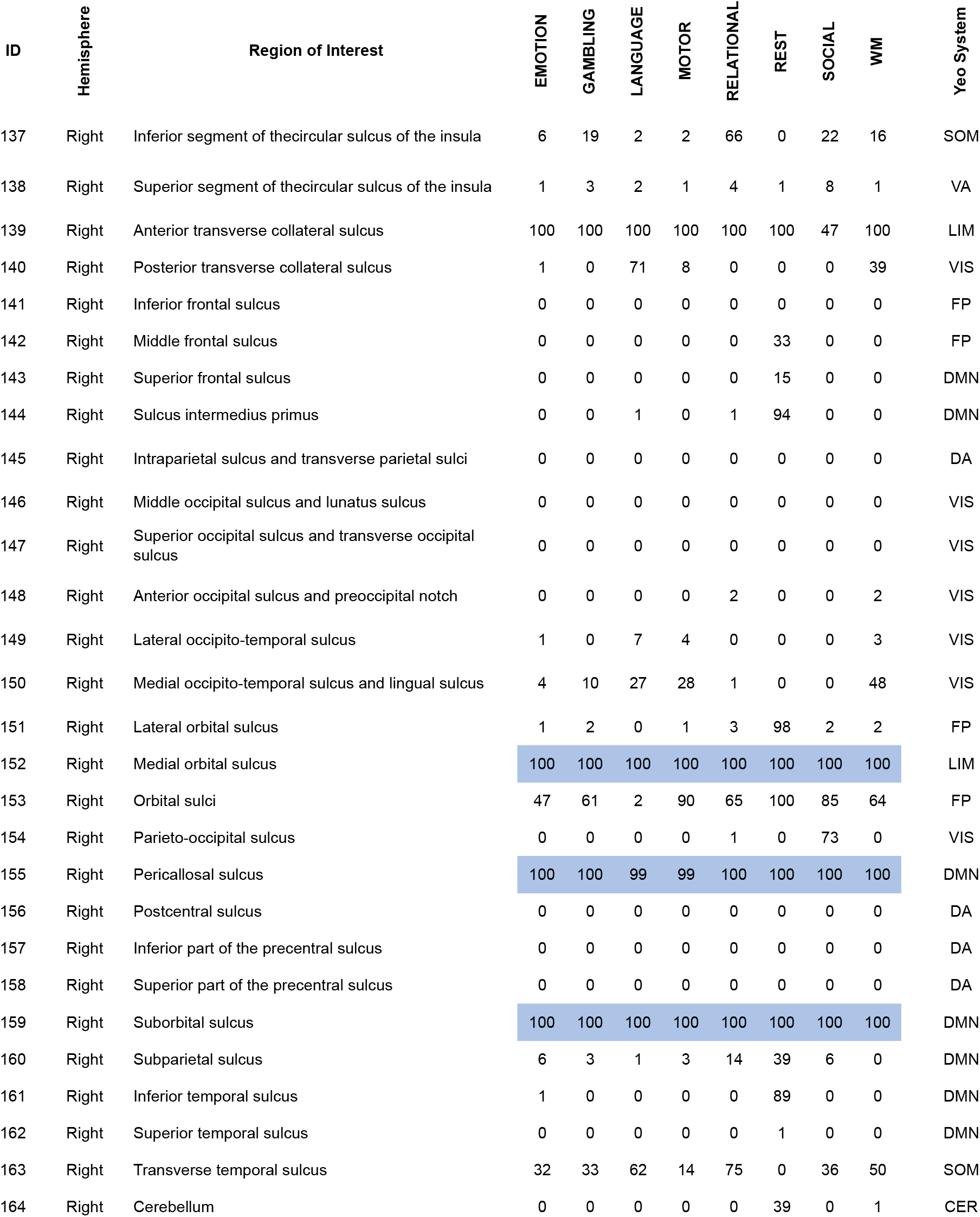

## References

[1] E. Bullmore and O. Sporns, “Complex brain networks: graph theoretical analysis of structural and functional systems,” Nature reviews neuroscience, vol. 10, no. 3, pp. 186–198, 2009.

[2] D. S. Bassett and O. Sporns, “Network neuroscience,” Nature neuroscience, vol. 20, no. 3, p. 353, 2017.

[3] S. Mori and J. Zhang, “Principles of diffusion tensor imaging and its applications to basic neuroscience research,” Neuron, vol. 51, no. 5, pp. 527–539, 2006.

[4] O. Sporns, G. Tononi, and R. Kötter, “The human connectome: a structural description of the human brain,” PLoS computational biology, vol. 1, no. 4, 2005.

[5] P. Hagmann, L. Cammoun, X. Gigandet, R. Meuli, C. J. Honey, V. J. Wedeen, and O. Sporns, “Mapping the structural core of human cerebral cortex,” PLoS biology, vol. 6, no. 7, 2008.

[6] S. Ogawa, T.-M. Lee, A. R. Kay, and D. W. Tank, “Brain magnetic resonance imaging with contrast dependent on blood oxygenation,” proceedings of the National Academy of Sciences, vol. 87, no. 24, pp. 9868–9872, 1990.

[7] M. W. Cole, D. S. Bassett, J. D. Power, T. S. Braver, and S. E. Petersen, “Intrinsic and task-evoked network architectures of the human brain,” Neuron, vol. 83, no. 1, pp. 238–251, 2014.

[8] C. J. Honey, J.-P. Thivierge, and O. Sporns, “Can structure predict function in the human brain?,” Neuroimage, vol. 52, no. 3, pp. 766–776, 2010.

[9] S. Smith, “Linking cognition to brain connectivity,” Nature neuroscience, vol. 19, no. 1, p. 7, 2016.

[10] K. Batista-García-Ramó and C. I. Fernández-Verdecia, “What we know about the brain structure–function relationship,” Behavioral Sciences, vol. 8, no. 4, p. 39, 2018.

[11] L. E. Suárez, R. D. Markello, R. F. Betzel, and B. Misic, “Linking structure and function in macroscale brain networks,” Trends in Cognitive Sciences, 2020.

[12] M. G. Preti and D. Van De Ville, “Decoupling of brain function from structure reveals regional behavioral specialization in humans,” Nature communications, vol. 10, no. 1, pp. 1–7, 2019.

[13] P. Tewarie, B. Prasse, J. M. Meier, F. A. Santos, L. Douw, M. Schoonheim, C. J. Stam, P. Van Mieghem, and A. Hillebrand, “Mapping functional brain networks from the structural connectome: relating the series expansion and eigenmode approaches,” NeuroImage, p. 116805, 2020.

[14] A. Avena-Koenigsberger, B. Misic, and O. Sporns, “Communication dynamics in complex brain networks,” Nature Reviews Neuroscience, vol. 19, no. 1, p. 17, 2018.

[15] F. Abdelnour, H. U. Voss, and A. Raj, “Network diffusion accurately models the relationship between structural and functional brain connectivity networks,” Neuroimage, vol. 90, pp. 335–347, 2014.

[16] J. Goñi, M. P. van den Heuvel, A. Avena-Koenigsberger, N. V. de Mendizabal, R. F. Betzel, A. Griffa, P. Hagmann, B. Corominas-Murtra, J.-P. Thiran, and O. Sporns, “Resting-brain functional connectivity predicted by analytic measures of network communication,” Proceedings of the National Academy of Sciences, vol. 111, no. 2, pp. 833–838, 2014.

[17] B. Mišić, R. F. Betzel, A. Nematzadeh, J. Goni, A. Griffa, P. Hagmann, A. Flammini, Y.-Y. Ahn, and O. Sporns, “Cooperative and competitive spreading dynamics on the human connectome,” Neuron, vol. 86, no. 6, pp. 1518–1529, 2015.

[18] S. Gu, F. Pasqualetti, M. Cieslak, Q. K. Telesford, B. Y. Alfred, A. E. Kahn, J. D. Medaglia, J. M. Vettel, M. B. Miller, S. T. Grafton, et al., “Controllability of structural brain networks,” Nature communications, vol. 6, p. 8414, 2015.

[19] S. Gu, R. F. Betzel, M. G. Mattar, M. Cieslak, P. R. Delio, S. T. Grafton, F. Pasqualetti, and D. S. Bassett, “Optimal trajectories of brain state transitions,” NeuroImage, vol. 148, pp. 305–317, 2017.

[20] J. D. Medaglia, “Clarifying cognitive control and the controllable connectome,” Wiley Interdisciplinary Reviews: Cognitive Science, vol. 10, no. 1, p. e1471, 2019.

[21] Y.-Y. Liu, J.-J. Slotine, and A.-L. Barabási, “Controllability of complex networks,” nature, vol. 473, no. 7346, pp. 167–173, 2011.

[22] C. Tu, R. P. Rocha, M. Corbetta, S. Zampieri, M. Zorzi, and S. Suweis, “Warnings and caveats in brain controllability,” NeuroImage, vol. 176, pp. 83–91, 2018.

[23] F. Pasqualetti, S. Gu, and D. S. Bassett, “Re: Warnings and caveats in brain controllability,” NeuroImage, vol. 197, pp. 586–588, 2019.

[24] N. U. Dosenbach, D. A. Fair, F. M. Miezin, A. L. Cohen, K. K. Wenger, R. A. Dosenbach, M. D. Fox, A. Z. Snyder, J. L. Vincent, M. E. Raichle, et al., “Distinct brain networks for adaptive and stable task control in humans,” Proceedings of the National Academy of Sciences, vol. 104, no. 26, pp. 11073–11078, 2007.

[25] J. D. Power and S. E. Petersen, “Control-related systems in the human brain,” Current opinion in neurobiology, vol. 23, no. 2, pp. 223–228, 2013.

[26] M. Omrani, C. D. Murnaghan, J. A. Pruszynski, and S. H. Scott, “Distributed task-specific processing of somatosensory feedback for voluntary motor control,” Elife, vol. 5, p. e13141, 2016.

[27] B. R. Eisenreich, R. Akaishi, and B. Y. Hayden, “Control without controllers: toward a distributed neuroscience of executive control,” Journal of cognitive neuroscience, vol. 29, no. 10, pp. 1684–1698, 2017.

[28] F. Pasqualetti, S. Zampieri, and F. Bullo, “Controllability metrics, limitations and algorithms for complex networks,” IEEE Transactions on Control of Network Systems, vol. 1, no. 1, pp. 40–52, 2014.

[29] D. C. Van Essen, S. M. Smith, D. M. Barch, T. E. Behrens, E. Yacoub, K. Ugurbil, W.-M. H. Consortium, et al., “The wu-minn human connectome project: an overview,” Neuroimage, vol. 80, pp. 62–79, 2013.

[30] T. M. Karrer, J. Z. Kim, J. Stiso, A. E. Kahn, F. Pasqualetti, U. Habel, and D. S. Bassett, “A practical guide to methodological considerations in the controllability of structural brain networks,” Journal of Neural Engineering, vol. 17, no. 2, p. 026031, 2020.

[31] E. S. Finn, X. Shen, D. Scheinost, M. D. Rosenberg, J. Huang, M. M. Chun, X. Papademetris, and R. T. Constable, “Functional connectome fingerprinting: identifying individuals using patterns of brain connectivity,” Nature neuroscience, vol. 18, no. 11, p. 1664, 2015.

[32] U. Tipnis, E. Amico, M. Ventresca, and J. Goni, “Modeling communication processes in the human connectome through cooperative learning,” IEEE Transactions on Network Science and Engineering, 2018.

[33] G. Deco, V. K. Jirsa, and A. R. McIntosh, “Emerging concepts for the dynamical organization of resting-state activity in the brain,” Nature Reviews Neuroscience, vol. 12, no. 1, pp. 43–56, 2011.

[34] C. Destrieux, B. Fischl, A. Dale, and E. Halgren, “Automatic parcellation of human cortical gyri and sulci using standard anatomical nomenclature,” Neuroimage, vol. 53, no. 1, pp. 1–15, 2010.

[35] B. Thomas Yeo, F. M. Krienen, J. Sepulcre, M. R. Sabuncu, D. Lashkari, M. Hollinshead, J. L. Roffman, J. W. Smoller, L. Zöllei, J. R. Polimeni, et al., “The organization of the human cerebral cortex estimated by intrinsic functional connectivity,” Journal of neurophysiology, vol. 106, no. 3, pp. 1125–1165, 2011.

[36] B. Mišić, R. F. Betzel, M. A. De Reus, M. P. Van Den Heuvel, M. G. Berman, A. R. McIntosh, and O. Sporns, “Network-level structure-function relationships in human neocortex,” Cerebral Cortex, vol. 26, no. 7, pp. 3285–3296, 2016.

[37] Y. Osmanlıoğlu, B. Tunç, D. Parker, M. A. Elliott, G. L. Baum, R. Ciric, T. D. Satterthwaite, R. E. Gur, R. C. Gur, and R. Verma, “System-level matching of structural and functional connectomes in the human brain,” Neuroimage, vol. 199, pp. 93–104, 2019.

[38] R. G. Bettinardi, G. Deco, V. M. Karlaftis, T. J. Van Hartevelt, H. M. Fernandes, Z. Kourtzi, M. L. Kringelbach, and G. Zamora-López, “How structure sculpts function: unveiling the contribution of anatomical connectivity to the brain’s spontaneous correlation structure,” Chaos: An Interdisciplinary Journal of Nonlinear Science, vol. 27, no. 4, p. 047409, 2017.

[39] C. Seguin, M. P. Van Den Heuvel, and A. Zalesky, “Navigation of brain networks,” Proceedings of the National Academy of Sciences, vol. 115, no. 24, pp. 6297–6302, 2018.

[40] A. Avena-Koenigsberger, X. Yan, A. Kolchinsky, M. van den Heuvel, P. Hagmann, and O. Sporns, “A spectrum of routing strategies for brain networks,” PLoS computational biology, vol. 15, no. 3, p. e1006833, 2019.

[41] R. Lambiotte, R. Sinatra, J.-C. Delvenne, T. S. Evans, M. Barahona, and V. Latora, “Flow graphs: Interweaving dynamics and structure,” Physical Review E, vol. 84, no. 1, p. 017102, 2011.

[42] M. Breakspear, “Dynamic models of large-scale brain activity,” Nature neuroscience, vol. 20, no. 3, p. 340, 2017.

[43] R.F. Galán, “On how network architecture determines the dominant patterns of spontaneous neural activity,” PloS one, vol. 3, no. 5, 2008.

[44] S. El Boustani and A. Destexhe, “A master equation formalism for macroscopic modeling of asynchronous irregular activity states,” Neural computation, vol. 21, no. 1, pp. 46–100, 2009.

[45] M. T. Schaub, Y. N. Billeh, C. A. Anastassiou, C. Koch, and M. Barahona, “Emergence of slow-switching assemblies in structured neuronal networks,” PLoS computational biology, vol. 11, no. 7, 2015.

[46] C. E. Sexton, K. B. Walhovd, A. B. Storsve, C. K. Tamnes, L. T. Westlye, H. Johansen-Berg, and A. M. Fjell, “Accelerated changes in white matter microstructure during aging: a longitudinal diffusion tensor imaging study,” Journal of Neuroscience, vol. 34, no. 46, pp. 15425–15436, 2014.

[47] E. Tang, C. Giusti, G. L. Baum, S. Gu, E. Pollock, A. E. Kahn, D. R. Roalf, T. M. Moore, K. Ruparel, R. C. Gur, et al., “Developmental increases in white matter network controllability support a growing diversity of brain dynamics,” Nature communications, vol. 8, no. 1, pp. 1–16, 2017.

[48] B. A. Sauerbrei, J.-Z. Guo, J. D. Cohen, M. Mischiati, W. Guo, M. Kabra, N. Verma, B. Mensh, K. Branson, and A. W. Hantman, “Cortical pattern generation during dexterous movement is input-driven,” Nature, vol. 577, no. 7790, pp. 386–391, 2020.

[49] E. Amico, A. Arenas, and J. Goñi, “Centralized and distributed cognitive task processing in the human connectome,” Network Neuroscience, vol. 3, no. 2, pp. 455–474, 2019.

[50] P. T. Bell and J. M. Shine, “Subcortical contributions to large-scale network communication,” Neuroscience & Biobehavioral Reviews, vol. 71, pp. 313–322, 2016.

[51] R. Shadmehr and J. W. Krakauer, “A computational neuroanatomy for motor control,” Experimental brain research, vol. 185, no. 3, pp. 359–381, 2008.

[52] D. Ketteler, F. Kastrau, R. Vohn, and W. Huber, “The subcortical role of language processing. high level linguistic features such as ambiguity-resolution and the human brain; an fmri study,” NeuroImage, vol. 39, no. 4, pp. 2002–2009, 2008.

[53] M. R. Delgado, L. E. Nystrom, C. Fissell, D. Noll, and J. A. Fiez, “Tracking the hemodynamic responses to reward and punishment in the striatum,” Journal of neurophysiology, vol. 84, no. 6, pp. 3072–3077, 2000.

[54] L. F. Koziol and D. E. Budding, Subcortical structures and cognition: Implications for neuropsychological assessment. Springer Science & Business Media, 2009.

[55] G. E. Alexander, M. R. DeLong, and P. L. Strick, “Parallel organization of functionally segregated circuits linking basal ganglia and cortex,” Annual review of neuroscience, vol. 9, no. 1, pp. 357–381, 1986.

[56] M. W. Cole, J. R. Reynolds, J. D. Power, G. Repovs, A. Anticevic, and T. S. Braver, “Multi-task connectivity reveals flexible hubs for adaptive task control,” Nature neuroscience, vol. 16, no. 9, p. 1348, 2013.

[57] M. D. Greicius, B. Krasnow, A. L. Reiss, and V. Menon, “Functional connectivity in the resting brain: a network analysis of the default mode hypothesis,” Proceedings of the National Academy of Sciences, vol. 100, no. 1, pp. 253–258, 2003.

[58] M. P. Van Den Heuvel and H. E. H. Pol, “Exploring the brain network: a review on resting-state fmri functional connectivity,” European neuropsychopharmacology, vol. 20, no. 8, pp. 519–534, 2010.

[59] T. Ito, K. R. Kulkarni, D. H. Schultz, R. D. Mill, R. H. Chen, L. I. Solomyak, and M. W. Cole, “Cognitive task information is transferred between brain regions via resting-state network topology,” Nature communications, vol. 8, no. 1, pp. 1–14, 2017.

[60] D. C. Van Essen, K. Ugurbil, E. Auerbach, D. Barch, T. Behrens, R. Bucholz, A. Chang, L. Chen, M. Corbetta, S. W. Curtiss, et al., “The human connectome project: a data acquisition perspective,” Neuroimage, vol. 62, no. 4, pp. 2222–2231, 2012.

[61] M. F. Glasser, S. N. Sotiropoulos, J. A. Wilson, T. S. Coalson, B. Fischl, J. L. Andersson, J. Xu, S. Jbabdi, M. Webster, J. R. Polimeni, et al., “The minimal preprocessing pipelines for the human connectome project,” Neuroimage, vol. 80, pp. 105–124, 2013.

[62] G. Rosenthal, F. Váša, A. Griffa, P. Hagmann, E. Amico, J. Goñi, G. Avidan, and O. Sporns, “Mapping higher-order relations between brain structure and function with embedded vector representations of connectomes,” Nature communications, vol. 9, no. 1, p. 2178, 2018.

[63] M. Jenkinson, C. F. Beckmann, T. E. Behrens, M. W. Woolrich, and S. M. Smith, “Fsl,” Neuroimage, vol. 62, no. 2, pp. 782–790, 2012.

[64] J.-D. Tournier, R. Smith, D. Raffelt, R. Tabbara, T. Dhollander, M. Pietsch, D. Christiaens, B. Jeurissen, C.-H. Yeh, and A. Connelly, “Mrtrix3: A fast, flexible and open software framework for medical image processing and visualisation,” NeuroImage, p. 116137, 2019.

[65] R. E. Smith, J.-D. Tournier, F. Calamante, and A. Connelly, “Anatomically-constrained tractography: improved diffusion mri streamlines tractography through effective use of anatomical information,” Neuroimage, vol. 62, no. 3, pp. 1924–1938, 2012.

[66] B. Jeurissen, J.-D. Tournier, T. Dhollander, A. Connelly, and J. Sijbers, “Multi-tissue constrained spherical deconvolution for improved analysis of multi-shell diffusion mri data,” NeuroImage, vol. 103, pp. 411–426, 2014.

[67] J. D. Tournier, F. Calamante, and A. Connelly, “Improved probabilistic streamlines tractography by 2nd order integration over fibre orientation distributions,” in Proceedings of the international society for magnetic resonance in medicine, vol. 18, p. 1670, Ismrm, 2010.

[68] R. E. Smith, J.-D. Tournier, F. Calamante, and A. Connelly, “Sift2: Enabling dense quantitative assessment of brain white matter connectivity using streamlines tractography,” Neuroimage, vol. 119, pp. 338–351, 2015.

[69] J. D. Power, A. Mitra, T. O. Laumann, A. Z. Snyder, B. L. Schlaggar, and S. E. Petersen, “Methods to detect, characterize, and remove motion artifact in resting state fmri,” Neuroimage, vol. 84, pp. 320–341, 2014.

[70] D. Marcus, J. Harwell, T. Olsen, M. Hodge, M. Glasser, F. Prior, M. Jenkinson, T. Laumann, S. Curtiss, and D. Van Essen, “Informatics and data mining tools and strategies for the human connectome project,” Frontiers in neuroinformatics, vol. 5, p. 4, 2011.

[71] J. Z. Kim, J. M. Soffer, A. E. Kahn, J. M. Vettel, F. Pasqualetti, and D. S. Bassett, “Role of graph architecture in controlling dynamical networks with applications to neural systems,” Nature physics, vol. 14, no. 1, p. 91, 2018.

[72] B. Mohar, Y. Alavi, G. Chartrand, and O. Oellermann, “The laplacian spectrum of graphs,” Graph theory, combinatorics, and applications, vol. 2, no. 871–898, p. 12, 1991.

[73] L. A. Wolsey and G. L. Nemhauser, Integer and combinatorial optimization, vol. 55. John Wiley & Sons, 1999.

[74] L. v. d. Maaten and G. Hinton, “Visualizing data using t-sne,” Journal of machine learning research, vol. 9, no. Nov, pp. 2579–2605, 2008.

